# Gucy1α1 specifically marks kidney, heart, lung and liver fibroblasts

**DOI:** 10.1101/2024.05.15.594404

**Authors:** Valeria Rudman-Melnick, Davy Vanhoutte, Kaitlynn Stowers, Michelle Sargent, Mike Adam, Qing Ma, Anne Karina T. Perl, Alexander G. Miethke, Ashley Burg, Tiffany Shi, David A. Hildeman, E. Steve S. Woodle, J. Matthew Kofron, Prasad Devarajan

## Abstract

Fibrosis is a common outcome of numerous pathologies, including chronic kidney disease (CKD), a progressive renal function deterioration. Current approaches to target activated fibroblasts, key effector contributors to fibrotic tissue remodeling, lack specificity. Here, we report Gucy1α1 as a specific kidney fibroblast marker. Gucy1α1 levels significantly increased over the course of two clinically relevant murine CKD models and directly correlated with established fibrosis markers. Immunofluorescent (IF) imaging showed that Gucy1α1 comprehensively labelled cortical and medullary quiescent and activated fibroblasts in the control kidney and throughout injury progression, respectively. Unlike traditionally used markers platelet derived growth factor receptor beta (Pdgfrβ) and vimentin (Vim), Gucy1α1 did not overlap with off-target populations such as podocytes. Notably, Gucy1α1 labelled kidney fibroblasts in both male and female mice. Furthermore, we observed elevated GUCY1α1 expression in the human fibrotic kidney and lung. Studies in the murine models of cardiac and liver fibrosis revealed Gucy1α1 elevation in activated Pdgfrβ-, Vim- and alpha smooth muscle actin (αSma)-expressing fibroblasts paralleling injury progression and resolution. Overall, we demonstrate Gucy1α1 as an exclusive fibroblast marker in both sexes. Due to its multiorgan translational potential, GUCY1α1 might provide a novel promising strategy to specifically target and mechanistically examine fibroblasts.

## Introduction

Fibrosis, defined as excessive extracellular matrix (ECM) deposition in interstitial spaces, is a final common outcome of numerous pathologies in different organs including kidney (1). Initially a crucial part of tissue repair, progressive fibrosis results in maladaptive organ remodeling in many chronic human conditions, such as CKD (2). The extent of interstitial fibrotic remodeling serves as a powerful predictor of overall adverse outcomes and inversely correlates with estimated glomerular filtration rate (eGFR) which is a measure of kidney filtration capacity and CKD progression (3–5). According to the United States Renal Data System annual report for 2022, the percentage of US adults with CKD is 14%, reaching 33.2% among those aged ≥65 years (6). Advanced CKD also might transition towards end-stage kidney disease (ESKD), a debilitating condition which results in ultimate kidney failure and death (7). With kidney replacement therapy remaining merely supportive and transplant options limited, targeted intervention strategies for CKD will require advanced mechanistic knowledge of the cellular and molecular processes that drive fibrosis (8). Moreover, dissecting the mechanisms of pathologic fibrotic tissue remodeling may eventually help to treat chronic pathologies in other organs such as heart, lung and liver, which could occur concurrently or independent of kidney dysfunction (9–11).

Fibroblasts are key cellular contributors to the fibrotic response and normal parenchymal loss observed in CKD (12). Under physiologic conditions, spindle-shaped fibroblasts reside in the interstitial space of the kidney, where they help to ensure tissue architecture and homeostasis. However, disproportionate or repetitive acute kidney injury (AKI) results in fibroblast activation, excessive proliferation, and ECM overproduction (13), which disrupts normal kidney structure and lead to organ failure. Some fibroblasts express contractile genes, such as *Acta2* (encoding αSma), and are therefore called “myofibroblasts” (14). Recent single-cell RNA-sequencing (scRNA-seq) analysis of human CKD demonstrated that activated fibroblasts exhibit highest ECM related gene expression scores, indicating their role as main ECM-producing cells in the chronically diseased kidney (15). Thus, targeting fibroblasts as the kidney cell population responsible for most injury-induced ECM deposition might represent a promising strategy to intercept maladaptive fibrotic kidney remodeling. However, this remains challenging due to genetic and functional heterogeneity of fibroblasts and the lack of unique markers that that would allow for their comprehensive labeling with no off-target effects in other kidney cell populations (16).

The protein encoded by *Gucy1α1* (Gucy1α1 a.k.a. Gucy1α3) is an alpha subunit of soluble guanylate cyclase 1 (SG-1), a heterodimeric protein which is a primary receptor of nitric oxide (NO) (17). Genome-wide association studies (GWAS), and next-generation sequencing along with genetic and pharmacological targeting identified *Gucy1α1* among the genes influencing blood flow parameters and risks of cardiovascular diseases (18–23). Moreover, pharmacological stimulation of NO-SG-1-cGMP pathway alleviated kidney fibrosis and improved kidney function in rat models of kidney disease (24–26). According to the GUDMAP Consortium *in situ* hybridization data, embryonic murine kidney exhibits *Gucy1α1* expression in the cortical and medullary interstitium but not in the developing renal corpuscle or capillary loop (27, 28). scRNA-seq profiling also demonstrated that *Gucy1α1* labels several murine embryonic stromal populations, including nephron progenitor, cortical, medullary, collecting duct associated and ureteric stroma (29). In addition, an independent study revealed that *GUCY1Α1* is present in the human fetal kidney *COL1A1/2*-positive interstitial clusters with no expression in epithelial, endothelial or podocyte populations (30). However, Gucy1α1 potential as an adult kidney fibroblast marker remains poorly explored. Our scRNA-seq (31) demonstrated that *Gucy1α1* expression is restricted to three adult murine fibroblast populations and is reflective of fibrosis degree in advanced kidney injury. To further explore the potential of Gucy1α1 as a novel kidney fibroblast marker, we examined Gucy1α1 spatiotemporal expression changes over the course of murine CKD induced by unilateral ischemia-reperfusion (UIR) and ureter obstruction (UUO) (32–35). With a comprehensive array of validations, we revealed that Gucy1α1 labelled kidney cortical and medullary fibroblasts, including fractions expressing αSma, Pdgfrβ and Vim, in the control kidney and throughout injury progression. However, unlike these classically used markers, Gucy1α1 labelled fibroblasts both thoroughly and selectively, without overlapping with other populations such as podocytes. We also demonstrated the potential of Gucy1α1 as a kidney fibroblast marker in both sexes. Moreover, we observed GUCY1α1 expression in αSMA- and VIM-positive fibroblasts in the human kidney and lung. Finally, we unraveled multiorgan potential of Gucy1α1 as a fibroblast marker in the murine models of cardiac and liver fibrosis.

## Results

### *Gucy1α1* selectively marks kidney fibroblasts in two independent clinically relevant fibrosis models

We examined single-cell specific gene expression signatures of kidney fibroblasts in the clinically relevant models of advanced CKD induced by UIR and UUO (Supplemental Figure 1A). Our recent report (31) revealed that both control and prolonged fibrotic injuries exhibited three separate fibroblast clusters among other kidney cell populations, designated as “Fibro 1”, “Fibro 2” and “Fibro 3” due to their respective “secretory”, “contractile” and “migratory” transcriptional enrichments (Supplemental Figure 1B). Here, we found that *Gucy1α1* specifically marked all three fibroblast clusters. Unlike traditionally used stromal markers which only labelled some populations (*Postn, Col1α1, Pdgfrα, Acta2, Dcn, Des*) or overlapped with epithelial, endothelial and immune clusters (*Vim, Meis1/2, Lgals1, Tagln2*), *Gucy1α1* was enriched exclusively in the fibroblasts (Supplemental Figure 1, A and B). Using IF staining we validated our scRNA-seq findings and showed Gucy1α1 expression in the control cortical and medullary kidney stroma, which was markedly elevated in both our CKD models (Supplemental Figure 1C).

### Gucy1α1 levels increase as kidney injury progresses and correlate with key fibrosis markers

Next, we examined the spatiotemporal expression of Gucy1α1 in UIR and UUO, two models of AKI-to-CKD transition (36). Quantitative histological and molecular analysis at multiple timepoints following UIR/UUO (Day 1, 4, 7, 14 and 28) confirmed progressive increase in cortical and medullary ECM deposition along with crucial kidney fibroblast markers, such as Pdgfrβ, αSma and Vim (Figure 1, A-E, Supplemental figure 2, A-D and 3, A-D). Thus, our models elicited key features of progressive CKD, including fibroblast activation, “contractile” αSma-positive phenotype acquisition and ECM overproduction, resulting in maladaptive kidney fibrotic remodeling (37, 38). Then we used our models to examine Gucy1α1 expression over the course of kidney fibrosis progression. We found that *Gucy1α1* mRNA and protein levels increased as both injuries progressed, reaching significance at Day 4 in UUO (Figure 2, A-C, Supplemental figures 4A and 5A). Furthermore, we demonstrated a significant correlation between Gucy1α1 and established fibroblast markers Pdgfrβ, αSma and Vim in both UIR and UUO models (Figure 2D and E). Of note, Gucy1α1 correlated particularly significantly with αSma, a smooth muscle cell marker indicating contractile phenotype in fibroblasts (16). Collectively, we showed that progressive Gucy1α1 elevation accompanied AKI-to-CKD transition and correlated with crucial historical markers of kidney fibrosis.

**Figure 1.**
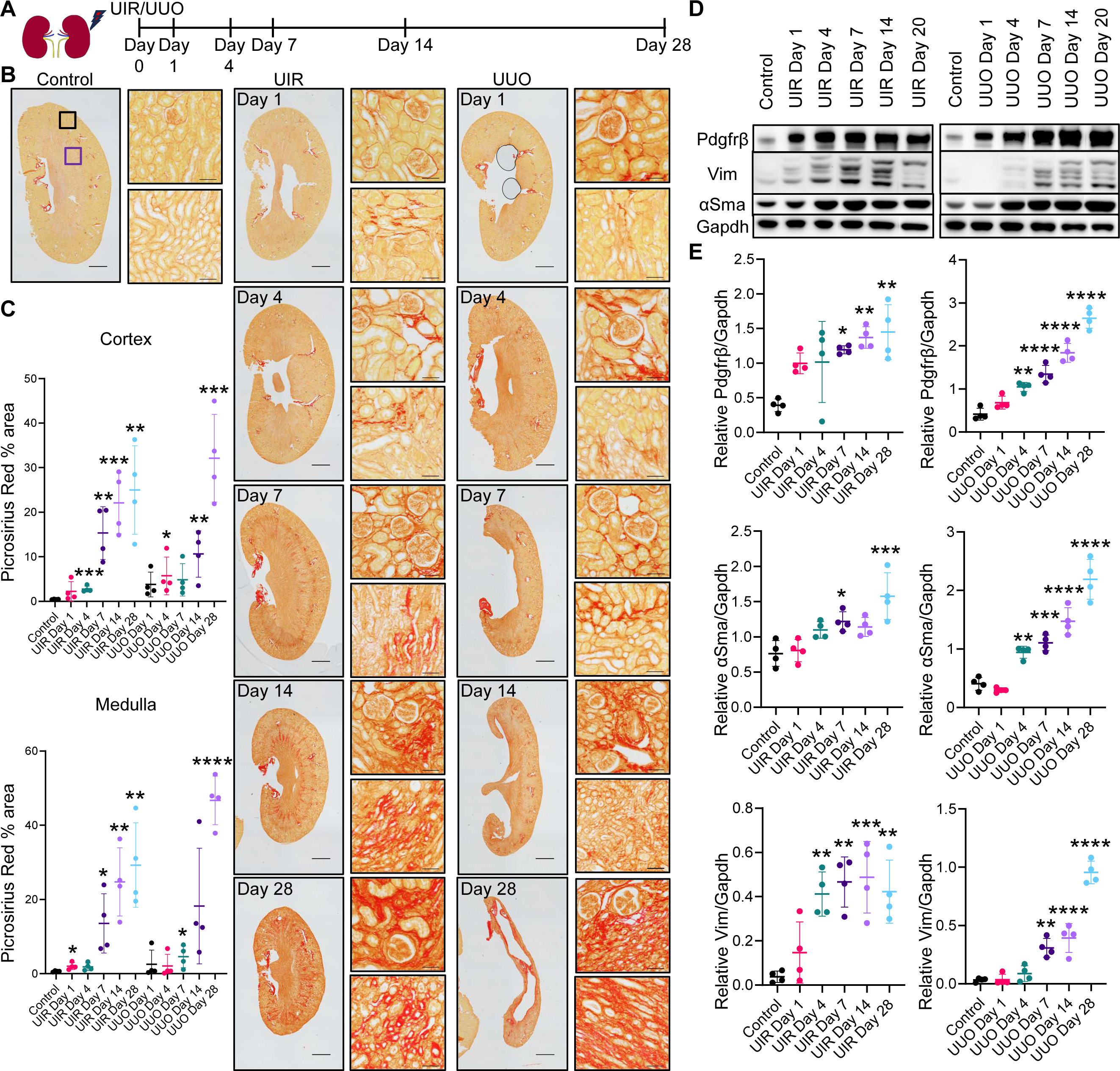
UIR and UUO induced murine CKD models exhibited progressive kidney fibrotic remodeling. (**A**) Schematic of the murine CKD progression model experimental timeline. (**B** and **C**) Picrosirius Red staining showing ECM accumulation over the course of UIR and UUO. (**B**) Representative whole kidney images (original magnification, ×4, 1.48 μm/px zoom, scale 1000 μm) and ×20 0.30 μm/px zoom (cortical – upper images, exemplifying location highlighted with black frames on the control kidney image, medullary – lower images, purple frames, scale 50 μm). (**C**) Quantification of total staining in kidney cortex and medulla, n=4 per group, unpaired 2-tailed *t*-test, P *≤0.05, **≤0.01, ***≤0.001, ****≤0.0001 compared to the control. (**D** and **E**) Western blotting showing gradual fibrosis markers upregulation over the course of UIR and UUO. (**D**) Representative Pdgfrβ, αSma, Vim, Gapdh bands. (**E**) Western blotting quantification, Pdgfrβ, αSma, Vim signal normalized to Gapdh, n=4 per group, ordinary one-way ANOVA, P *≤0.05, **≤0.01, ***≤0.001, ****≤0.0001 compared to the control. Data in scatter plots is presented as mean ± SD.

**Figure 2.**
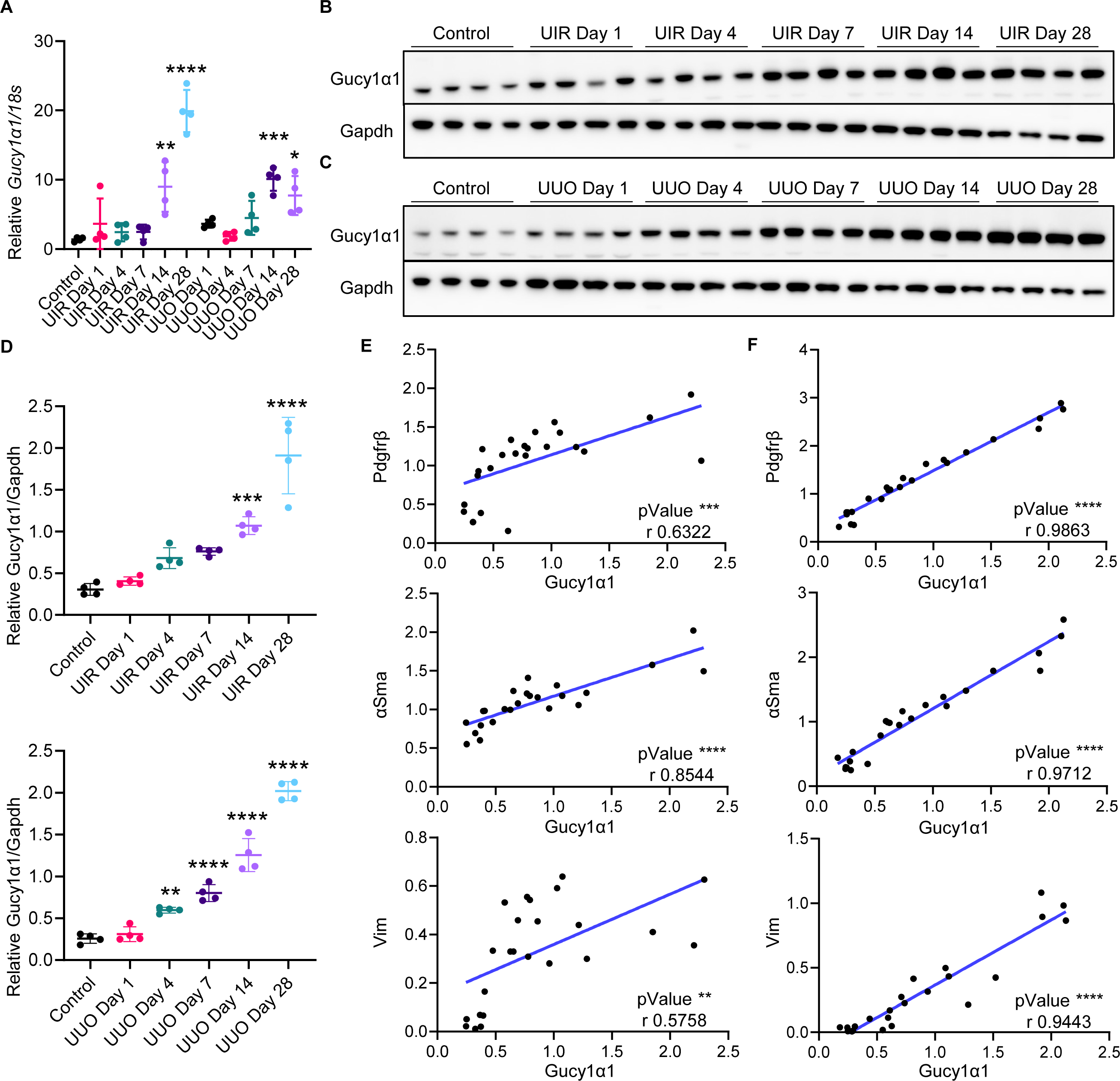
Gucy1α1 levels increase as kidney injury progresses and correlate with key fibrosis markers. (**A**) qPCR showing *Gucy1α1* expression changes over the course of UIR and UUO. Relative expression normalized to *18s*, shown as fold change, n=4 per group, ordinary one-way ANOVA, P *≤0.05, **≤0.01, ***≤0.001, ****≤0.0001 compared to the control. (**B-D**) Western blotting demonstrating progressive Gucy1α1 elevation accompanying fibrosis progression in both models, images (**B** and **C**) and quantification (**D**). Gucy1α1 signal normalized to Gapdh, n=4 per group, ordinary one-way ANOVA, P **≤0.01, ***≤0.001, ****≤0.0001 compared to the control. (**E** and **F**) Correlation analysis between normalized Pdgfrβ, αSma, Vim levels and Gucy1α1 in the control and UIR (**E**) and control and UUO (**F**) models at all experimental timepoints. Pearson r correlation analysis, n=24 per marker (control, Day 1, Day 4, Day 7, Day 14, Day 28, n=4 per each group), r values and P as **≤0.01, ***≤0.001, ****≤0.0001 for each pair are shown on the graphs presenting simple linear regression of correlation between Pdgfrβ, αSma, Vim vs Gucy1α1. Data in scatter plots is presented as mean ± SD.

### Gucy1α1 comprehensively labels cortical and medullary kidney fibroblasts

Next, we examined spatial localization of Gucy1α1 over the course of UIR and UUO induced kidney fibrosis. To account for distinctive phenotypical features and origins and to further dissect regional heterogeneity (39), we separately characterized cortical and medullary kidney fibroblasts. Using high-resolution multichannel IF imaging, we revealed moderate Gucy1α1 expression in the control cortical interstitium, which demonstrated progressive increase as the fibrotic remodeling progressed (Figure 3, A and B, Supplemental Figure 6, A-D). IF also showed that cortical Pdgfrβ, αSma and Vim protein levels progressively increased in the UIR/UUO treated kidneys compared to the control. Next, we used spatial quantitative IF analysis to dissect the molecular signature of Gucy1α1-positive fibroblasts and found near-total overlap between cortical Gucy1α1 and Pdgfrβ in the control and UIR/UUO treated kidneys (Figure 3, B and C, Supplemental Figure 6, E and F). We also found that while only a fraction of control cortical Gucy1α1-expressing fibroblasts was Vim-positive, more of them acquired persistent Vim expression over the course of both injuries (Figure 3, B and C, Supplemental Figure 6G). Most control cortical Gucy1α1-positive fibroblasts were negative for αSma, indicating their inactivated phenotype. However, Gucy1α1/αSma colocalization steadily increased along with fibrosis progression, reaching peak levels at UIR Day 7 and UUO Day 4 (Figure 3, B and C, Supplemental Figure 6H). Analogous to the cortex, UIR and UUO also caused significantly increased medullary Gucy1α1, Pdgfrβ, αSma and Vim protein levels compared to the control (Supplemental Figure 7, A-D). Furthermore, we observed up to 100% overlap between medullary Gucy1α1- and Pdgfrβ-expressing areas in the control kidney and throughout both injuries (Figure 4, A-C, Supplemental Figure 7, E and F). Of note, while cortical Gucy1α1-positive fibroblasts acquired Vim and αSma co-expression more progressively, Gucy1α1/αSma and Gucy1α1/Vim double positivity in the medulla increased abruptly at Day 1 and remained elevated throughout the course of both injuries (Figure 4, A-C, Supplemental Figure 7, G and H). Overall, these data demonstrate that Gucy1α1 comprehensively marks cortical and medullary fibroblasts in the normal kidney and at multiple stages of fibrosis progression, including in the αSma-positive fraction.

**Figure 3.**
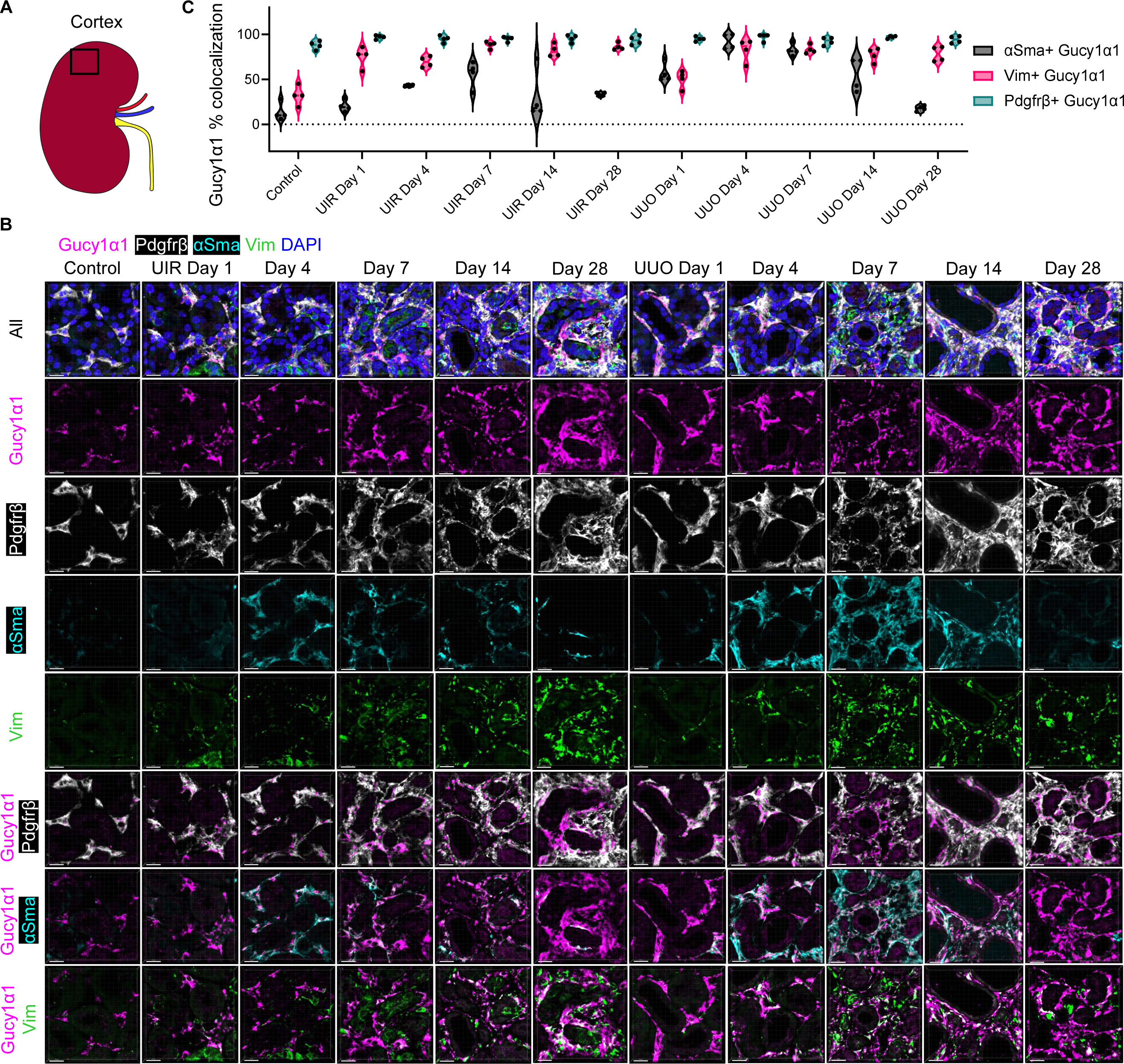
Gucy1α1 co-labels Pdgfrβ-, αSma- and Vim-positive fractions of the baseline and activated kidney fibroblasts in the cortical areas. (**A**) Schematic of the kidney anatomy with respect to cortical area (highlighted with a black frame). (**B**) Representative IF images of the control, UIR/UUO Day 1, 4, 7, 14 and 28 kidneys. Note remarkable degree of colocalization between Gucy1α1 (magenta) and Pdgfrβ (white). Also note that portions of Gucy1α1 expressing areas are positive for αSma (cyan) and Vim (green). DAPI, blue. The upper panels show all combined signals, the middle panels show individual channels, and three bottom panels demonstrate Gucy1α1 along with Pdgfrβ, αSma or Vim, respectively. Original magnification, × 60, maximal intensity projection from a Z-stack, 0.09 μm/px Nyquist zoom, scale 15 μm. (**C**) Violin plot showing quantitative IF analysis of cortical patterns of kidney fibrosis markers expression. Note near-total overlap between cortical Gucy1α1- and Pdgfrβ-positive areas (average percentage of Pdgfrβ co-expression in Gucy1α1-positive areas for control: 88%; UIR Day 1: 97%, Day 4: 95%, Day 7: 95%, Day 14: 95.5%, Day 28: 93.5%; UUO Day 1: 95%, Day 4: 97%, Day 7: 92.5%, Day 14: 97%, Day 28: 94.25%). On the contrary, only 13.8% of control cortical Gucy1α1-positive interstitial areas exhibited αSma positivity. This number rose as both injuries progressed, peaked at UIR Day 7 (average 56%) and UUO Day 4 (average 92.25%) and then declined. Most cortical Gucy1α1-expressing areas retained Vim co-expression throughout both injuries. N=4 animals per group. Only interstitial non-glomerular areas were included in the analysis.

**Figure 4.**
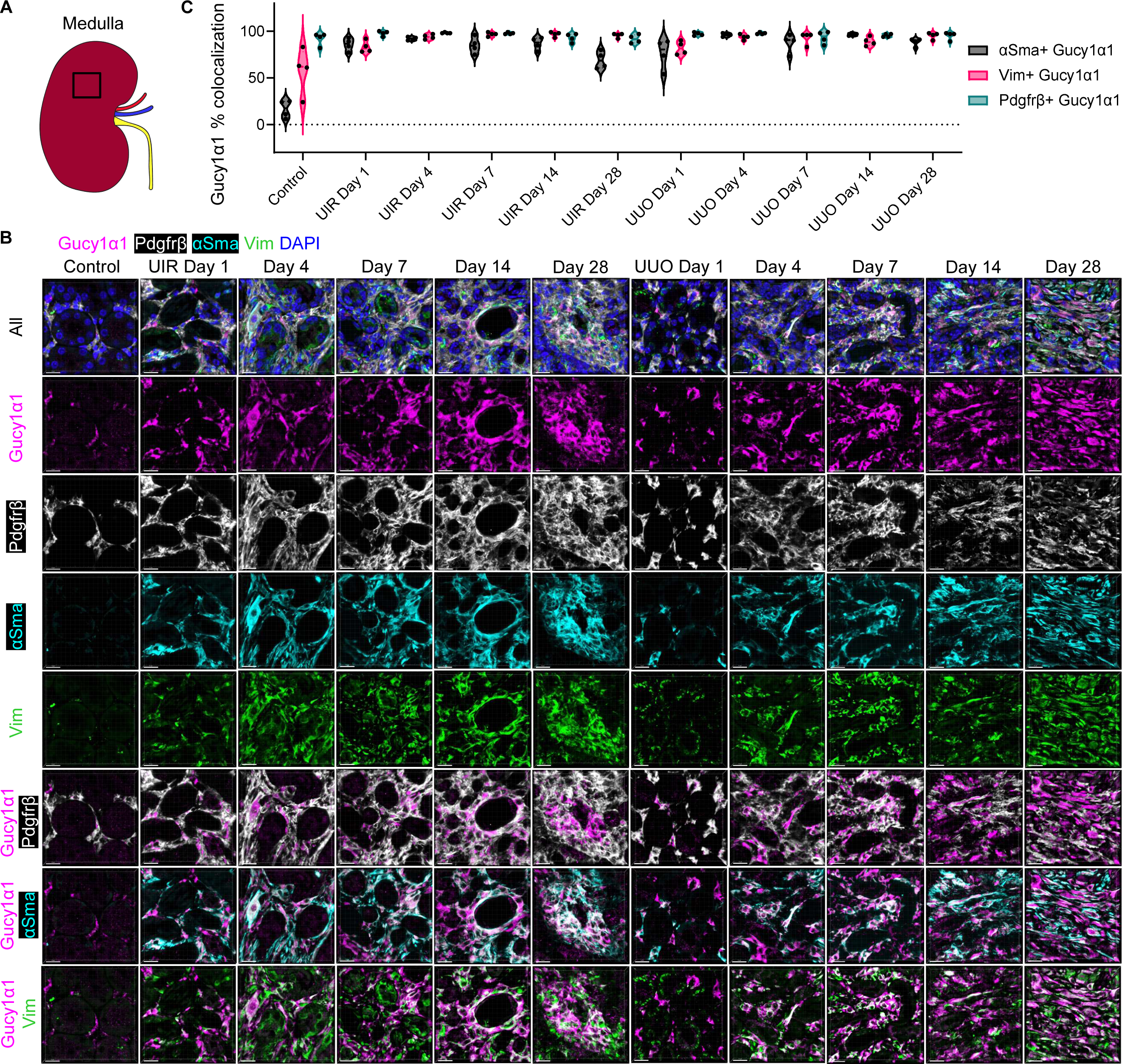
Gucy1α1 co-labels Pdgfrβ-, αSma- and Vim-positive fractions of the baseline and activated kidney fibroblasts in the medullary areas. (**A**) Schematic of the kidney anatomy with respect to medullary area (highlighted with a black frame). (**B**) Representative IF images of the control, UIR/UUO Day 1, 4, 7, 14 and 28 kidneys, Gucy1α1 (magenta), Pdgfrβ (white), αSma (cyan), Vim (green), DAPI, blue. The upper panels show all combined signals, the middle panels show individual channels, and three bottom panels demonstrate Gucy1α1 along with Pdgfrβ, αSma or Vim, respectively. Original magnification, × 60, maximal intensity projection from a Z-stack, 0.09 μm/px Nyquist zoom, scale 15 μm. (**C**) Violin plot showing quantitative IF analysis of medullary patterns of kidney fibrosis markers expression. Note near-total overlap between Gucy1α1- and Pdgfrβ-positive areas in the control kidneys and at all stages of both injuries (average percentage of Pdgfrβ co-expression in Gucy1α1-positive areas for control: 92%; UIR Day 1: 97.75%, Day 4: 98.25%, Day 7: 97.75%, Day 14: 93%, Day 28: 91.75%; UUO Day 1: 97%, Day 4: 97.75%, Day 7: 93.25%, Day 14: 95.5%, Day 28: 95.25%). Also note that medullary Gucy1α1-expressing fibroblasts acquired αSma co-expression more abruptly and robustly than the cortical ones, and retained it at higher percentages (average double positivity for control: 16.5%; UIR Day 1: 85.75%, Day 4: 92%, Day 7: 85%, Day 14: 86.75%, Day 28: 69.25%; UUO Day 1: 76%, Day 4: 96.25%, Day 7: 87.5%, Day 14: 96.5%, Day 28: 88.25%). Similar to the cortex, most medullary Gucy1α1-positive areas exhibited Vim co-expression. N=4 animals per group.

### Gucy1α1 specifically marks kidney fibroblasts throughout the course of UIR and UUO induced fibrosis

We next tested the specificity of Gucy1α1 as a novel kidney fibroblast marker. scRNA-seq predicted that *Gucy1α1* exclusively labels kidney fibroblasts under the normal conditions and in advanced injuries with no off-target expression in endothelial, immune, tubular or podocyte populations (Supplemental Figure 1, A and B). We sought to validate these findings on the protein level and at multiple stages of UIR and UUO induced kidney fibrosis. Particularly, we assessed intraglomerular Gucy1α1 expression compared to the established stromal markers Vim and Pdgfrβ. We found that both Vim and Pdgfrβ exhibited significantly higher baseline intraglomerular expression and colocalized with nephrin (Nphs1) positive podocytes, in comparison to Gucy1α1 which was not expressed in podocytes (Figure 5, A-C). This pattern remained constant throughout the course of both injuries, with Gucy1α1 only expressed in the periglomerular stroma. Collectively, we demonstrated that Gucy1α1 comprehensively and selectively labels kidney fibroblasts throughout the course of murine CKD progression.

**Figure 5.**
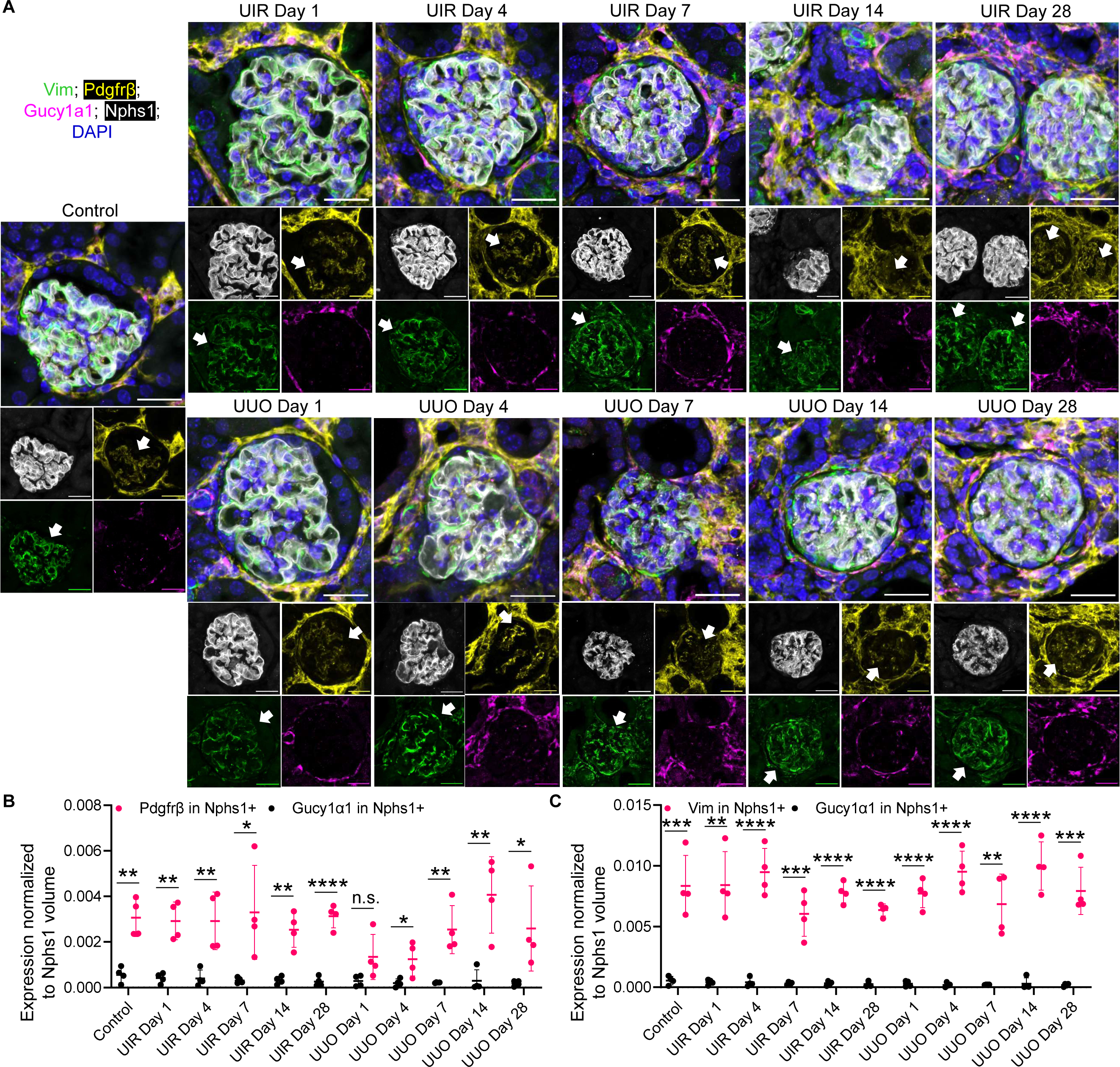
Gucy1α1 does not label glomerular populations compared to historically used kidney fibrosis markers. (**A**) Representative IF images showing Gucy1α1 (magenta), Pdgfrβ (yellow) and Vim (green), Nphs1 (white), DAPI, blue. Pdgfrβ and Vim are abundantly present inside the glomerulus, including colocalization with Nphs1-positive podocytes (shown with white arrows). Original magnification, × 60, maximal intensity projection from a Z-stack, 0.08 μm/px Nyquist zoom, scale 25 μm (**B** and **C**) Quantitative IF analysis of intraglomerular Gucy1α1, Pdgfrβ and Vim expression in the control and UIR/UUO treated kidneys. Gucy1α1, Pdgfrβ and Vim signals are normalized to Nphs1-expressing area volume and averaged from all the glomeruli captured in the imaging field, n=4 animals per group. Gucy1α1 and Pdgfrβ (**B**) or Gucy1α1 and Vim (**C**) normalized averaged intraglomerular signals are compared using multiple unpaired *t*-test with FDR correction for multiple comparisons. P values are shown as P *≤0.05, **≤0.01, ***≤0.001, ****≤0.0001, n.s., not significant between each pair of markers at each timepoint. (**B**) q values for Gucy1α1 vs Pdgfrβ comparisons: control: 0.001729; UIR Day 1: 0.001729, Day 4: 0.006212, Day 7: 0.017478, Day 14: 0.001729, Day 28: 0.000237; UUO Day 1: 0.037973, Day 4: 0.018007, Day 7: 0.004218, Day 14: 0.004218, Day 28: 0.021214. (**C**) q values for Gucy1α1 vs Vim comparisons: control: 0.001093; UIR Day 1: 0.001279, Day 4: 0.000177, Day 7: 0.001093, Day 14: 0.000013, Day 28: 0.000005; UUO Day 1: 0.000059, Day 4: 0.000102, Day 7: 0.001838, Day 14: 0.000177, Day 28: 0.000327. Data in scatter plots is presented as mean ± SD.

### Gucy1α1 marks kidney fibroblasts in the female model of murine CKD

Due to the significance of kidney fibrotic disease for both sexes (40), we sought to investigate whether Gucy1α1 also labels kidney fibroblasts in the female CKD. For this purpose, we established a female model of UIR induced kidney fibrosis via prolonged ischemia and harvested the kidneys at 28 days after the procedure. To ensure injury induction, we analyzed the degrees of ECM deposition with Picrosirius Red staining and showed that 50- and 60-minute ischemia caused a statistically significant fibrotic response in both cortex and medulla (Figure 6, A and B). We then examined Gucy1α1 expression with respect to progressively increasing ischemia time and degree of fibrosis.

**Figure 6.**
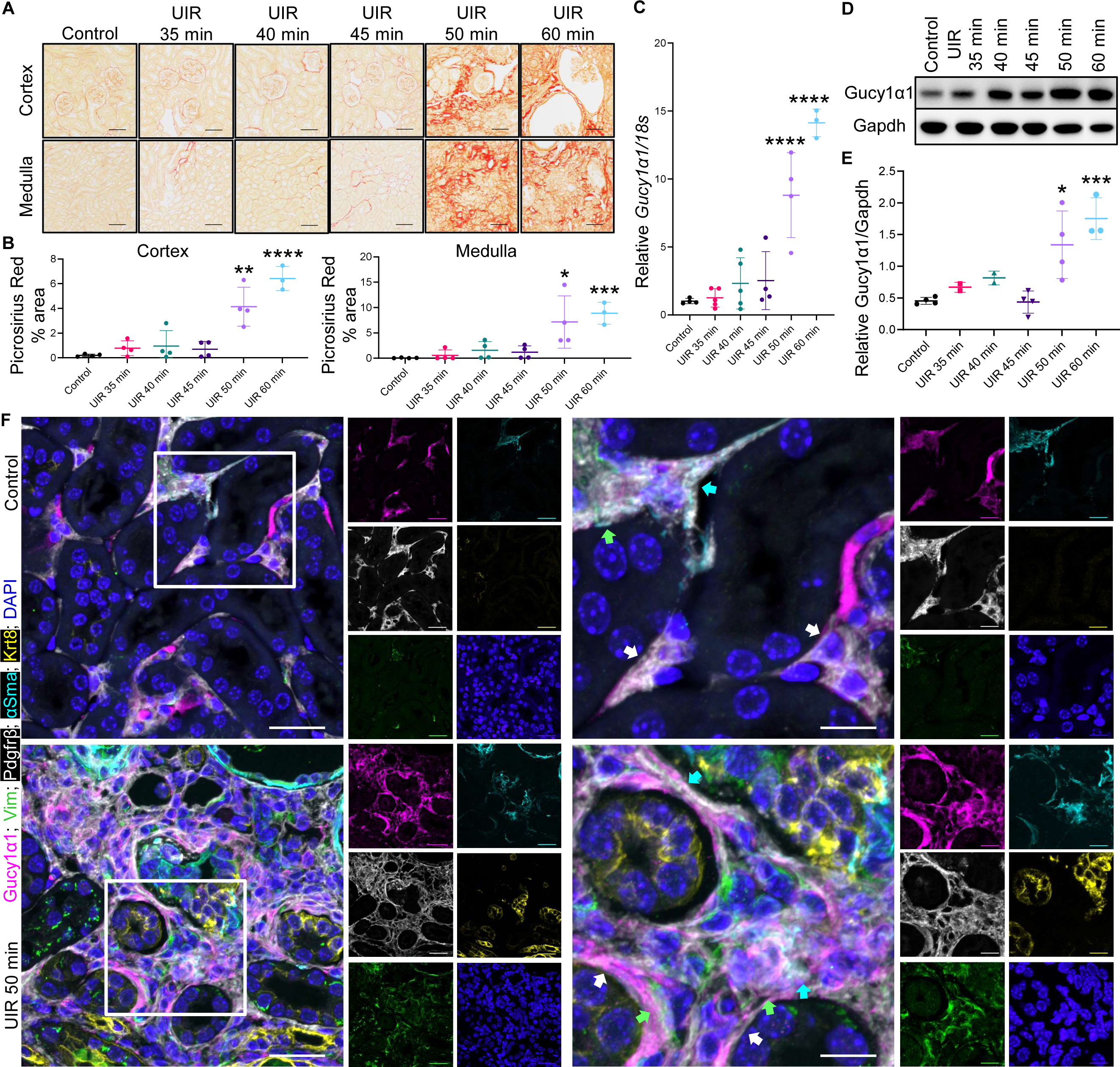
Gucy1α1 marks kidney fibroblasts in the female model of murine CKD. (**A** and **B**) Picrosirius Red representative images and quantification showing the effects of prolonged UIR on the female kidney. (**A**) Original magnification, ×20, 0.30 μm/px zoom, scale 50 μm. (**B**) Quantification of total staining in kidney cortex and medulla, n=3-5 per group, unpaired 2-tailed *t*-test, P *≤0.05, **≤0.01, ***≤0.001, ****≤0.0001 compared to the control. (**C**) qPCR showing *Gucy1α1* expression changes in the female CKD model. Relative expression normalized to *18s*, shown as fold change, n=3-5 per group, ordinary one-way ANOVA, ****≤0.0001 compared to the control. (**D** and **E**) Western blotting revealing that 50 and 60 min UIR caused significant Gucy1α1 upregulation in the female CKD model. Representative bands (**D**) and quantification (**E**), Gucy1α1 signal normalized to Gapdh, n=2-4 per group, ordinary one-way ANOVA, P *≤0.05, ***≤0.001 compared to the control. (**F**) IF images showing Gucy1α1 expression in the female normal and fibrotic kidneys. 50 min UIR caused remarkable stromal expansion, accompanied by fibrosis markers Gucy1α1 (magenta), Pdgfrβ (white), αSma (cyan) and Vim (green) elevation between Krt8 (yellow) positive injured epithelial tubules. Note colocalization between Gucy1α1 and Pdgfrβ (white arrows), αSma (cyan arrows) and Vim (green arrows) in both control and fibrotic conditions. DAPI, blue, original magnification, × 60, maximal intensity projection from a Z-stack, 0.14 μm/px Nyquist zoom, scale 25 μm (left side) and 0.06 μm/px Nyquist zoom, scale 10 μm (right side, highlighted with white frames). Data in scatter plots is presented as mean ± SD.

We found that *Gucy1α1* RNA and protein levels parallelled the severity of fibrotic remodeling, increasing significantly in 50- and 60-minute UIR compared to the control (Figure 6, C-E, Supplemental Figure 8, A and B). Spatial examination using multichannel IF demonstrated that Gucy1α1 was expressed in the interstitial spaces of normal kidney, labeling Pdgfrβ-, Vim- and αSma-positive fibroblasts (Figure 6F). Moreover, UIR caused remarkable expansion of Gucy1α1-expressing fibroblasts located between cytokeratin 8 (Krt8) positive injured tubules (41) of the female kidney. Like the male models of murine CKD, female UIR demonstrated that Gucy1α1 thoroughly covers Pdgfrβ-, Vim- and αSma-expressing fractions of kidney fibroblasts. Importantly, our imaging revealed that while Vim was also expressed in Krt8-positive tubular epithelial cells, Gucy1α1 was not.

### Gucy1α1 exhibits translational and multiorgan potential as a novel fibroblast marker

To explore whether Gucy1α1 might possess a translational potential, we examined its expression in human biopsy specimens from healthy and fibrotic kidneys. First, we verified the onset of fibrotic remodeling in the biopsy specimen from human ESKD compared to the healthy kidney using Picrosirius Red staining (Figure 7A). Then, we used multichannel IF to detect GUCY1α1 in the human kidney. We found minimal GUCY1α1 expression in the normal kidney, colocalizing with αSMA-expressing areas (Figure 7B). Human ESKD specimen, on the contrary, exhibited a remarkable increase in GUCY1α1 protein and the appearance of GUCY1α1-/VIM-along with GUCY1α1-/VIM-/αSMA-positive fibroblasts. Moreover, we demonstrated that GUCY1α1 also labels human fibroblasts in the normal and interstitial pulmonary fibrosis (IPF) lung specimens. Using Picrosirius Red staining, we confirmed that IPF elicited substantial fibrotic remodeling, including excessive ECM deposition altering normal pulmonary architecture (Figure 8A). IF revealed minimal GUCY1α1 expression in the VIM-positive stromal cells within alveolar walls of normal lung and substantial GUCY1α1 elevation caused by IPF (Figure 8B). High-resolution imaging showed that, like the kidney, human lung fibrosis resulted in significant stromal expansion, comprised of GUCY1α1-/VIM- and GUCY1α1-/VIM-/αSMA-expressing fibroblast fractions. Collectively, these data indicate that GUCY1α1 holds multiorgan translational potential.

**Figure 7.**
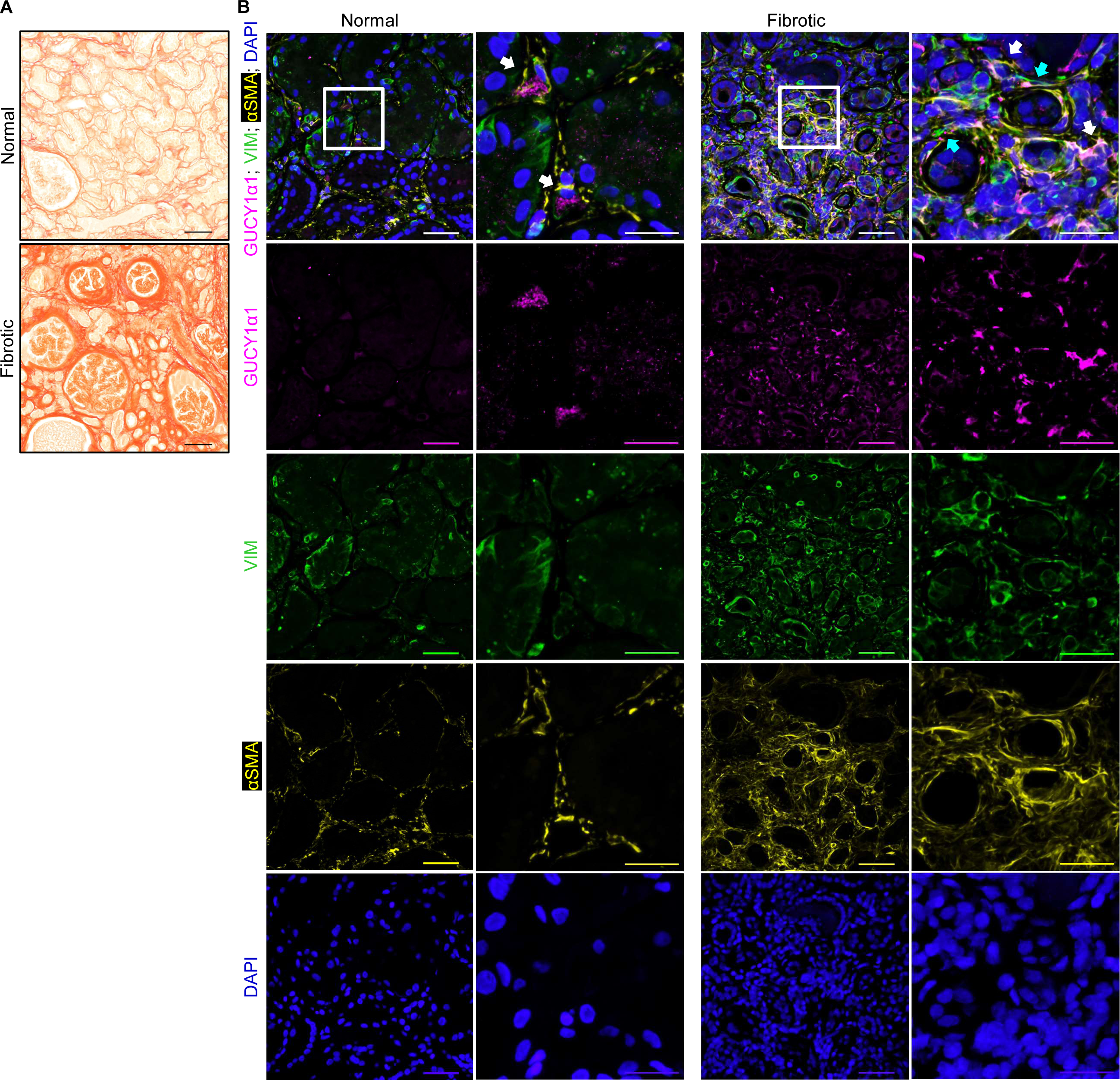
GUCY1α1 is expressed in the human kidney fibroblasts. (**A**) Picrosirius Red staining validating elevated ECM deposition in the fibrotic kidney compared to the normal control. Original magnification, ×20, 0.30 μm/px zoom, scale 100 μm. (**B**) IF showing mild interstitial GUCY1α1 (magenta) expression in the normal kidney, colocalized with αSMA (yellow). Fibrosis elicited GUCY1α1, αSMA and VIM (green) elevation. GUCY1α1 colocalization with other fibrosis markers is pointed out with white (αSMA) and blue (VIM) arrows. DAPI, blue, original magnification, × 60, maximal intensity projection from a Z-stack, 0.28 μm/px Nyquist resolution, scale 50 μm and 0.09 μm/px Nyquist zoom, scale 25 μm (highlighted with white frames).

**Figure 8.**
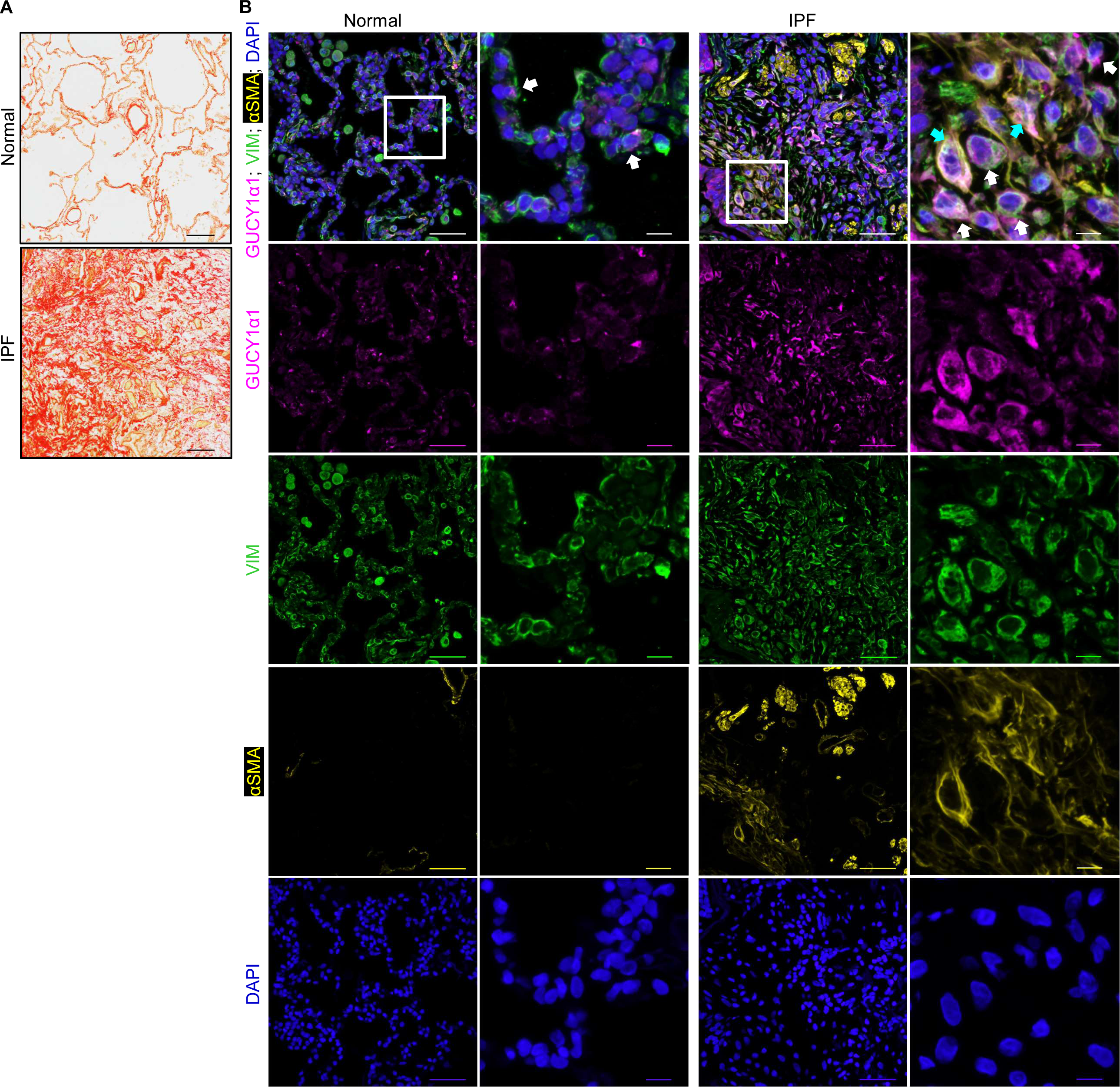
GUCY1α1 is expressed in the human lung fibroblasts. (**A**) Picrosirius Red staining validating robustly elevated ECM deposition in the IPF specimen compared to the normal control lung. Original magnification, ×20, 0.30 μm/px zoom, scale 100 μm. (**B**) IF showing moderate GUCY1α1 (magenta) expression in the normal lung, colocalized with VIM (green) expressing cells of alveolar walls (shown with white arrows). IPF elicited GUCY1α1, αSMA (yellow) and VIM elevation. Note that IPF elicited double GUCY1α1-/VIM-(pointed out with white arrows) and triple GUCY1α1-/VIM-/αSMA-positive (blue arrows) fibroblast fractions. DAPI, blue, original magnification, × 60, maximal intensity projection from a Z-stack, 0.28 μm/px Nyquist resolution, scale 50 μm and 0.08 μm/px Nyquist zoom, scale 10 μm (highlighted with white frames).

To further address multiorgan potential of Gucy1α1, we conducted additional studies using murine fibrosis models in other organs. Cardiac fibrosis induced by myocardial infarction (MI) (42) exhibited significant upregulation of classical stromal markers such as Pdgfrβ, Vim and αSma, which reached prominence at Day 3 and 7 post-MI (Figure 9, A-C, Supplemental Figure 9, A-E). Of note, Gucy1α1 levels reached a plateau at Day 7 and remained significantly elevated until Day 28 post-MI, thus coinciding with the fibrotic wound healing phase following MI (Figure 9, B-D). IF detected low interstitial expression of Pdgfrβ, Vim, αSma and Gucy1α1 in the normal myocardium, with minimal double Pdgfrβ-/Gucy1α1-positive fibroblast staining (Figure 9, E). However, MI caused pronounced fibrotic tissue expansion, evidenced by elevated interstitial Pdgfrβ, Vim, αSma and Gucy1α1 levels. High-resolution imaging revealed that Gucy1α1 was expressed in Pdgfrβ-, Vim- and αSma-positive fibroblast fractions at Day 28 post-MI.

**Figure 9.**
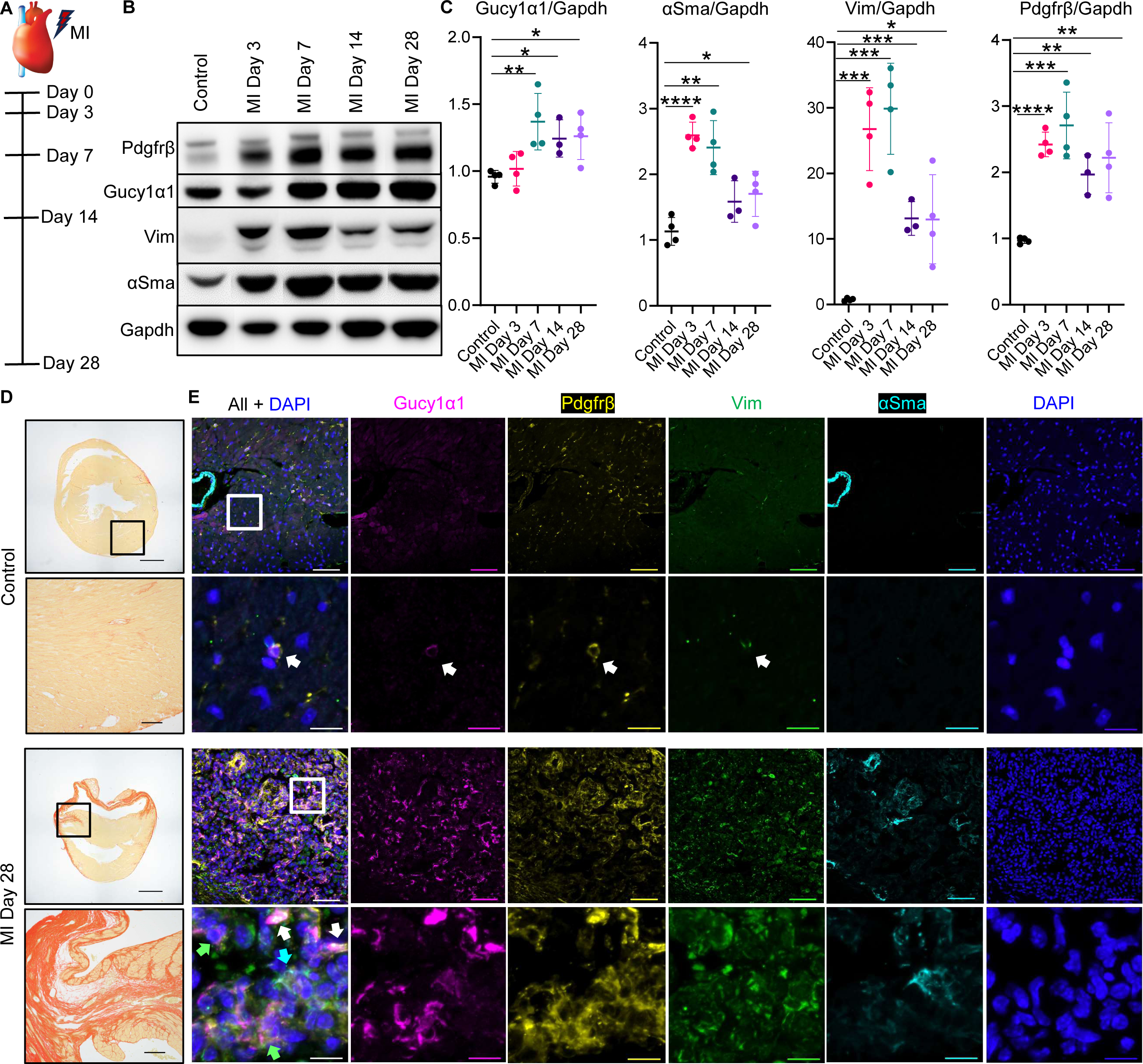
Gucy1α1 labels cardiac fibroblasts in the MI model of heart fibrosis. (**A**) Schematic of the MI progression model experimental timeline. (**B** and **C**) Western blotting showing Gucy1α1, Pdgfrβ, αSma and Vim expression changes over the course of MI. Representative bands (**B**) and quantification (**C**) of Gucy1α1, Pdgfrβ, αSma and Vim signal normalized to Gapdh. N=3-4 per group, unpaired 2-tailed *t*-test, P *≤0.05, **≤0.01, ***≤0.001, ****≤0.0001 compared to the control. Note that all fibrosis markers peaked at Day 3-7 post-MI. (**D**) Picrosirius Red staining demonstrating mature scar formation in the myocardial wall at MI Day 28 compared to the control heart. Whole heart image, original magnification, ×4, 1.48 μm/px zoom, scale 1000 μm and ×20 0.30 μm/px zoom, scale 100 μm (highlighted with black frames). (**E**) IF revealing Gucy1α1 expression in the normal and fibrotic heart. Gucy1α1 (magenta), Pdgfrβ (yellow), αSma (cyan), Vim (green) and DAPI (blue). Original magnification, ×60; upper panels - maximal intensity projection from a Z-stack, 0.28 μm/px Nyquist resolution, scale 50 μm; lower panels - 0.05 μm/px Nyquist zoom, scale 10 μm (highlighted with white frames). Note that in the normal heart Gucy1α1 exhibits episodic expression in Pdgfrβ-positive fibroblasts (white arrow). MI caused robust Gucy1α1 elevation and colocalization with Pdgfrβ-(white arrows), αSma-(cyan arrow) and Vim-positive (green arrow) areas. Data in scatter plots is presented as mean ± SD.

We also examined Gucy1α1 expression patterns in the setting of biliary fibrosis, induced by dietary introduction of 3,5-diethoxycarbonyl-1,4-dihydrocollidine (DDC) (43, 44). Using Picrosirius Red analysis, we demonstrated that 14 days of DDC dietary supplementation resulted in significant ECM accumulation which resolved 14 days following DDC withdrawal (Figure 10, A-C). We identified that baseline liver fibroblasts are positive for Pdgfrβ and Vim, with occasional αSma expression (Figure 10D). Gucy1α1 co-labelled some Pdgfrβ- and Vim-positive normal liver fibroblasts. However, 14 days of DDC treatment elicited significant elevation of Gucy1α1, Pdgfrβ, Vim and αSma protein levels, with remarkable colocalization between the classical fibroblast markers mentioned above and Gucy1α1 (Figure 10E). This elevation returned to the baseline levels as the injury resolved, with some remaining Gucy1α1 expression in Pdgfrβ- and Vim-positive liver fibroblasts. Moreover, we found that like the kidney, Gucy1α1 exhibited remarkable direct correlation with all three classical fibrosis markers, especially Vim and Pdgfrβ (Figure 10F, Supplemental Figure 10A). Overall, our studies have identified Gucy1α1 as a specific and multiorgan marker of murine and human fibroblasts.

**Figure 10.**
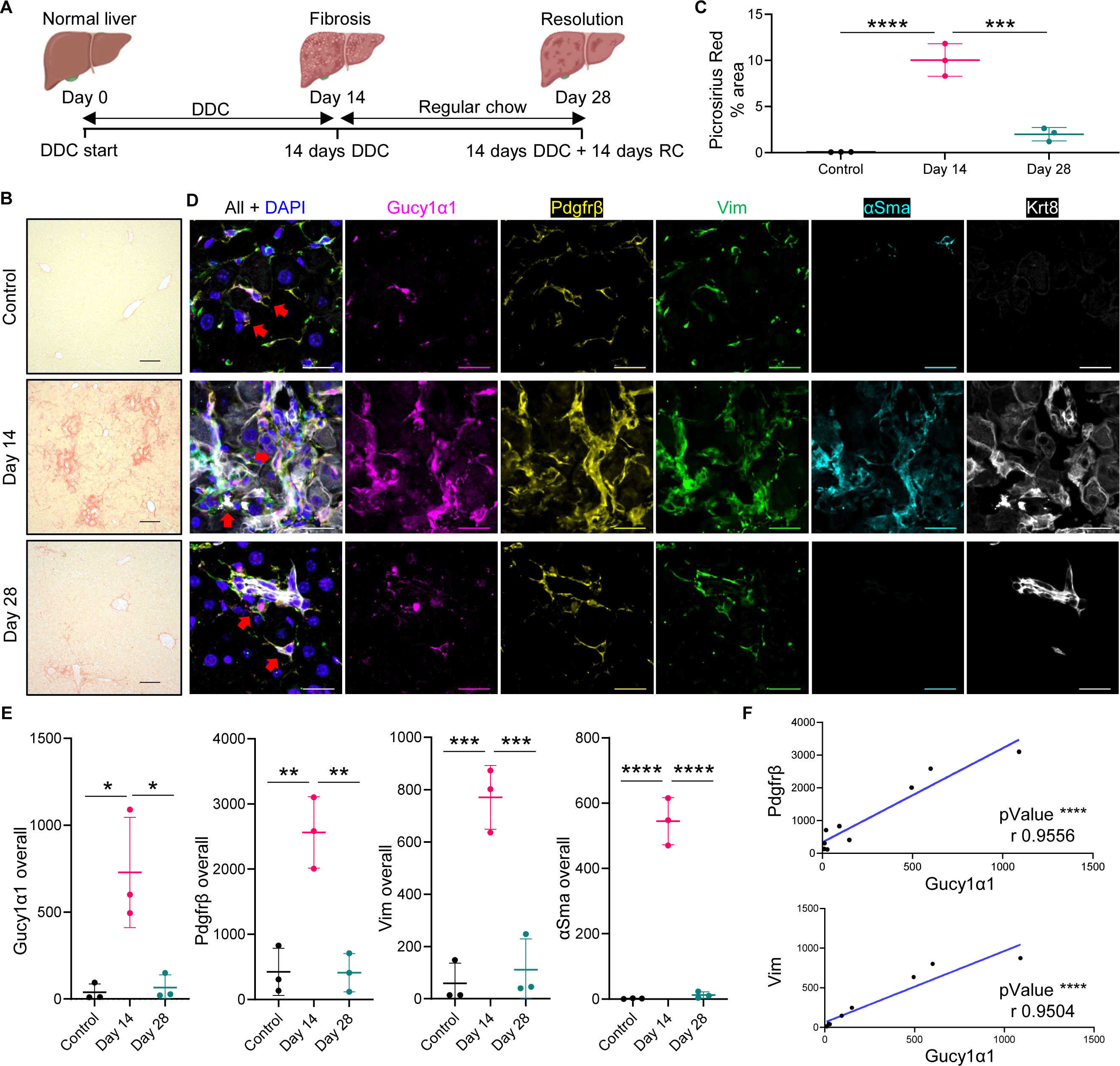
Gucy1α1 expression trajectory parallels DDC induced liver fibrosis resolution. (**A**) Schematic of the biliary fibrosis model. DDC is administered for 14 days, then regular chow (RC) is given. (**B** and **C**) Picrosirius Red staining. (**B**) Representative images, original magnification, ×20, 0.30 μm/px zoom, scale 100 μm. (**C**) Quantitative Picrosirius Red staining analysis, n=3 per group, ordinary one-way ANOVA, P ****≤0.0001 compared to the control. (**D** and **E**) IF demonstrating the expression of fibrosis markers in the liver over the course of DDC response. (**D**) Representative IF images demonstrate baseline Gucy1α1 (magenta) expression (red arrows) in the spindle-shaped Pdgfrβ-(yellow) and Vim-positive (green) liver fibroblasts. Baseline αSma (cyan) expression is very low. DDC administration results in dramatic Gucy1α1, Pdgfrβ, αSma and Vim upregulation and colocalization (areas of colocalization pointed with red arrows). 14 days after DDC withdrawal (Day 28) only traces of periductal (Krt8, white) and interstitial fibrosis remain (red arrows). DAPI, blue. Original magnification, ×60, maximal intensity projection from a Z-stack, 0.09 μm/px Nyquist resolution, scale 20 μm. (**E**) IF quantification, n=3 per group, ordinary one-way ANOVA, P **≤0.01, ***≤0.001, ****≤0.0001 compared to the control. (**F**) Correlation analysis between Pdgfrβ, Vim levels and Gucy1α1 at all the timepoints detected by IF. Pearson r correlation analysis, n=9 per marker (Control, Day 14, 28, n=3 per group), r values and P as ****≤0.0001 for each pair are shown on the graphs presenting simple linear regression of correlation between Pdgfrβ and Vim vs Gucy1α1. Data in scatter plots is presented as mean ± SD.

## Discussion

Developing a targeted approach to halt CKD progression remains challenging due to the complexity of molecular and cellular mechanisms underlying kidney fibrosis (45, 46). Transcriptional and functional heterogeneity of the key effector population contributing to aberrant ECM deposition and pathologic tissue remodeling – kidney fibroblasts – remains a crucial challenge in the field. While recent research has made significant progress in unraveling the molecular nature of fibrotic pathologies (2), a comprehensive strategy to exclusively trace and target kidney fibroblasts is still elusive, due to the lack of specificity of currently used markers (47–56). The current study presents Gucy1α1 as a newly validated kidney marker capable of labelling fibroblasts both comprehensively and selectively.

We and others recently used single-cell or single-nucleus RNA-seq to dissect the heterogeneity of kidney fibroblasts (15, 31, 57). Consistent with Li. et al. (57), we observed transcriptional and functional heterogeneity among kidney fibroblasts, which were divided based on their gene expression enrichments into “secretory” (“Fibro 1”), “contractile” (“Fibro 2”) and “migratory” (“Fibro 3”) clusters (31). *Gucy1α1* comprehensively marked secretory (high in ECM related genes such as *Col1a1/2, Col3a1, Fn1*), contractile (*Acta2, Myh11, Myl9* high) and migratory (*Sfrp2, Pdgfrβ, Amotl1* high) fibroblasts with no off-target expression in any other epithelial, endothelial, immune or podocyte populations. We therefore examined Gucy1α1 as a novel marker that might allow for tracing and targeting kidney fibroblasts in an exclusive manner at multiple AKI-to-CKD transition stages. With that goal, we established two independent clinically relevant murine models of kidney fibrosis caused by UIR or UUO (58) which both elicited progressive kidney fibrotic remodeling. We found that *Gucy1α1* mRNA and protein levels progressively increased as fibrosis in both models advanced. Moreover, we discovered a significant direct correlation between Gucy1α1 and classical fibrotic remodeling markers, such as Pdgfrβ, αSma and Vim. Pdgfrβ is traditionally used as a kidney fibroblast marker in many studies (12, 15). Mildly expressed in the normal kidney stroma, including glomerular mesangium and interstitial fibroblasts (59), Pdgfrβ becomes significantly elevated in both murine and human kidney fibrosis (60). Our previous study (31) showed that *Pdgfrβ* is present in all three fibroblast fractions at advanced stages of UIR and UUO. *Acta2*, which marks contractile “myofibroblast” phenotype, predominantly labelled “Fibro 2” populations of control and fibrotic kidneys with mild to absent expression in other fibroblast clusters. We also showed that *Vim*, while exhibiting medium expression levels in all three fibroblast populations, was also present in many off-target clusters, including immune cells and podocytes. In contrast, Gucy1α1 marked Vim- and αSma-expressing fibroblast fractions and exhibited near-total overlap with Pdgfrβ-positive interstitial fibroblasts in the control kidney and throughout all stages of fibrosis progression in both models. Since regional heterogeneity might represent a challenge to comprehensive kidney fibroblast labelling (39), we separately assessed Gucy1α1 potential in cortex and medulla and found that it equally comprehensively labels kidney fibroblasts in both regions. The trajectory of double Gucy1α1/αSma and Gucy1α1/Vim positivity might reflect the process of resident fibroblast activation and “contractile” phenotype acquisition by a portion of them. Our analysis identified a regional molecular heterogeneity inside kidney stroma, while only a fraction of cortical Gucy1α1-positive fibroblasts retained αSma expression throughout the whole course of both injuries, most medullary fibroblasts remained double Gucy1α1/αSma positive at all stages of UIR and UUO. This distinction might reflect regional heterogeneity in “contractile” phenotype acquisition by activated kidney fibroblasts. Also, a higher percentage of cortical and medullary fibroblasts from UUO treated kidneys exhibited double Gucy1α1/αSma positivity, probably reflecting the more severe fibrotic response caused by UUO compared to UIR (31, 57).

Notably, Gucy1α1 demonstrated no intraglomerular presence, compared to significant Pdgfrβ expression which colocalized with podocyte marker Nphs1, in both control and fibrotic kidneys. Previous research (61) also showed robust Pdgfrβ upregulation in the cells occupying glomerular Bowman’s space triggered by focal segmental glomerulosclerosis. Furthermore, recent scRNA-seq (62) identified *PDGFRΒ* expression in many glomerular cell types, including almost all human and some murine podocytes, murine parietal epithelial cells along with human and murine mesangial-like cells. Thus, we and others demonstrated that while Pdgfrβ labels interstitial fibroblasts, it also exhibits off-target expression among many glomerular populations. Collectively, these findings indicate that Gucy1α1 might serve as a better marker for exclusive labeling of all interstitial quiescent and activated fibroblasts with no off-target effects in the glomerulus.

Due to the increased importance of accounting for sex differences in the kidney molecular landscapes and injury response (63–65), we conducted a set of experiments to evaluate Gucy1α1 in the female murine CKD model. We found that *Gucy1α1* mRNA and protein levels reflected the degree of ECM deposition in the female UIR induced fibrosis. Female kidneys exhibited intratubular pattern of Gucy1α1 expression, which underwent robust upregulation following UIR and overlapped with Pdgfrβ-, αSma- and Vim-positive interstitial areas. These findings suggest that Gucy1α1 holds a potential as a specific fibroblast marker in both sexes.

Another challenge in the field is to identify reliable fibroblast labelling strategies that are applicable to both experimental models and human specimens with kidney fibrosis (16). Since Gucy1α1 levels are reflective of fibrosis degree in the murine UIR and UUO models, it might serve as a predictor of kidney injury progression trajectory either towards recovery or exacerbated fibrotic remodeling and CKD/ESKD. GUCY1α1 expression in αSMA- and VIM-expressing cells of the normal human kidney, and its upregulation in the fibrotic human kidney, suggests that it can be used as a marker of kidney fibroblasts and predictor of kidney injury outcome in humans. Our finding corroborated scRNA-seq and single nuclear RNA-seq (snRNA-seq) predicted *GUCY1Α1* expression in the human kidney stromal cells, such as fibroblasts, myofibroblasts and pericytes, reported by Kidney Precision Medicine Project (66). We also revealed mild GUCY1α1 expression in the normal human lung which was elevated in interstitial pulmonary fibrosis (IPF), colocalizing with αSMA- and VIM. This human finding is consistent with the previous murine scRNA-seq data, showing *Gucy1a1* expression in the pulmonary ECM-producing *Acta2*-positive populations (67).

Recent scRNA-seq analysis (68) identified *Gucy1a1* as a pericyte marker in lung and kidney. The authors defined pericytes based upon strong expression of *Cspg4* and *Pdgfrβ*. However, none of these genes is exclusive to pericytes. According to our scRNA-seq data (31), *Pdgfrβ* labels all three fibroblast populations including “contractile” *Acta2*-rich cluster. Likewise, *Cspg4* was present in the “contractile” fibroblasts along with Fibro3 “migratory” population. Thus, *Cspg4-*/*Pdgfrβ-*positive clusters identified as “pericyte-enriched” or even “stringent pericytes” might represent a subset of fibroblasts or fibroblast/pericyte mixture. To further address the multiorgan potential of Gucy1α1, we conducted a series of studies in other organs prone to fibrosis, such as heart and liver. In both organs, Gucy1α1 levels parallelled the trajectory of fibrotic injury progression and co-labelled baseline Pdgfrβ-/Vim-positive and activated Pdgfrβ-/Vim-/αSma-expressing fibroblasts. Our finding is consistent with the recent scRNA-seq which showed *Gucy1a1* enrichment in the portal fibroblasts and hepatic stellate cells in two other models of liver fibrosis (69). Overall, our findings highlight Gucy1α1 as a novel marker which selectively labelled kidney fibroblasts in both sexes and exhibited translational multiorgan potential.

## Methods

### Sex as a biological variable

For studies examining Gucy1α1 as a novel specific kidney fibroblast marker, both male and female models of CKD were used. To account for sex differences in response to the ischemic kidney injury (70), prolonged UIR was conducted in the female mice. Since Gucy1α1 was found to equally label male and female kidney fibroblasts, rest of the animal studies was performed in the male sex.

### Animal models

All animal experiments were performed in C57Bl/6 mice (Jackson Laboratory, #000664). <u>Kidney fibrosis</u> was induced in 10-week-old mice via atraumatic left renal pedicle clamping for 30 min at 37 °C (male UIR) or via irreversible left ureter ligation (male UUO). Kidneys were collected on Day 1, 4, 7, 14 and 28 (n=4 per group). Female mice were subjected to 35, 40, 45, 50 and 60 min UIR (n=3-5 per group for statistical analysis) and harvested at Day 28. Naïve mice of corresponding sexes (n=4) were used as controls. <u>Cardiac fibrosis</u> was induced by myocardial infarction (MI) as previously described (42). Mice were anesthetized using 2% isoflurane and a left lateral thoracotomy was performed. Next, the left coronary artery was identified and ligated with an 8-0 prolene suture (Cardinale Health, #8730H) just below the atrium. Sham-operated animals that were subjected to all procedures except permanent ligation were used as controls. Hearts were harvested at Day 3, 7, 14 and 28 post-MI and processed for further histological and molecular analysis (3-4 per group). <u>Biliary fibrosis</u> was induced in 8-week-old male C57Bl/6 mice via dietary supplementation of 5058 Lab diet chow admixed with 0.1% of DDC for up to 14 days to induce sclerosing cholangitis (43) and then changed to regular 5058 chow for another 14 days to model recovery from hepatobiliary injury. Groups of mice were harvested before (D0), during DDC treatment (D14), or after challenge (DDC D14+D14 regular diet) to obtain liver tissue samples for Picrosirius Red staining and IF (n=3 per group).

### Human tissue samples

Human normal paraffin kidney sections (male) were obtained from Novus Biologicals (T2234142). The fibrotic human kidney specimen was graciously donated by a deceased female with a prior medical history of ESKD and hypertension. Samples from this patient were obtained using a 16-gauge biopsy needle, placed in 10% formalin solution for 24 hours, then submitted to CCHMC Integrated Pathology Research Facility for paraffin-embedding. Deidentified human explant or donor lung tissues were processed at CCHMC Integrated Pathology Research Facility for paraffin-embedding.

### Microscopy and image analysis

Picrosirius Red staining was analyzed on BAF widefield Nikon NiE upright microscope. IF images were obtained on Nikon Ti-E AXR HD confocal microscope with the resonant scanner, processed with NIS-Elements AR 5.2.00 artificial intelligence denoise algorithm and analyzed with Bitplane Imaris 10.0.0. (71).

### Statistics

For the assessment of Gucy1α1 as a specific kidney fibroblast marker in the male CKD models, 4 animals per group were used across all the experiments. For the female CKD model, 3-5 animals per group were used for the statistical comparisons. For Gucy1α1 evaluation in the animal models of cardiac and liver fibrosis, 3-4 animals per group were used for all statistical comparisons. All the analysis was performed using GraphPad Prism 9. Comparisons between 2 unpaired, normally distributed data points were carried out via an unpaired 2-tailed *t* test. Comparisons between multiple groups were performed using one-way ANOVA with Tukey’s multiple-comparison test. For the analysis of glomerular Gucy1α1, Pdgfrβ and Vim expression, multiple unpaired *t* test with False Discovery Rate (FDR) correction for multiple comparisons was used. All data in scatter plots is presented as mean ± SD. Cortical and medullary analysis of colocalization between Gucy1α1, Pdgfrβ, αSma and Vim is shown as violin plots.

### Study approval

The Institutional Care and Use Committee (IACUC) of Cincinnati Children’s Hospital Medical Center approved all animal procedures in the study. All the experiments and methods, including animal husbandry and monitoring, were performed in accordance with relevant IACUC guidelines and regulations. Human kidneys donated for research were obtained from the local organ procurement office (LifeCenter) in accordance with institutional policies of Cincinnati Children’s Hospital Medical Center and the University of Cincinnati, and research protocols at LifeCenter.

<u>Detailed Methods</u> are available in the Supplement.

## Author contributions

VRM, DV, MA, KS, AKTP, AGM, DAH and PD conceived and designed the project, analyzed, and interpreted the data. VRM executed laboratory experiments, assembled the figures and prepared the manuscript. VRM and KS performed kidney injury modeling. MS performed heart injury modeling. DV collected heart samples for histological analysis and contributed to the corresponding Method section. KS and QM contributed to laboratory experiments and data analysis. MA provided bioinformatical data. JMK designed confocal imaging acquisition and analysis strategy and provided technical guidance. AKTP, AGM, AB, TS, DAH and ESSW provided human lung, murine liver, and human kidney samples, respectively, and contributed to the corresponding sections of the manuscript. VRM, DV, KS and PD performed final revisions of the manuscript. All authors read and approved the manuscript as submitted.

## Acknowledgements

This work was supported by NIH grants P50 DK096418, RO1 HL13395 and F30 AI167482.

