## Supplemental data for "Gucy1α1 specifically marks kidney, heart, lung and liver fibroblasts"

[1] Division of Nephrology and Hypertension, Cincinnati Children's Hospital Medical Center, Cincinnati, OH, USA. [2] Division of Molecular Cardiovascular Biology, Cincinnati Children's Hospital Medical Center, Cincinnati, OH, USA. [3] Division of Experimental Hematology and Cancer Biology, Cincinnati Children's Hospital Medical Center, Cincinnati, OH, USA. [4] Division of Neonatology and Pulmonary biology, Cincinnati Children's Hospital Medical Center, Cincinnati, OH, USA. [5] Division of Gastroenterology, Hepatology and Nutrition, Cincinnati Children's Hospital Medical Center, Cincinnati, OH, USA. [6] Division of Immunobiology, Cincinnati Children's Hospital Medical Center, Cincinnati, OH, USA. [7] Department of Surgery, University of Cincinnati College of Medicine, Cincinnati, OH, USA. [8] Division of Developmental Biology, Cincinnati Children's Hospital Medical Center, Cincinnati, OH, USA.

**Supplemental figures**

Figure S1. *Gucy1α1* selectively marks kidney fibroblasts in two independent clinically relevant fibrosis models.

Figure S2. UIR causes progressive elevation of fibrosis markers Vim, αSma and Pdgfrβ over the course of injury progression.

Figure S3. UUO causes progressive elevation of fibrosis markers Vim, αSma and Pdgfrβ over the course of injury progression.

Figure S4. UIR causes progressive elevation of Gucy1α1 over the course of injury progression.

Figure S5. UUO causes progressive elevation of Gucy1α1 over the course of injury progression.

Figure S6. Gucy1α1 co-labels Pdgfrβ-, Vim- and αSma-positive fractions of kidney fibroblasts in the cortical areas under the baseline conditions and over the course of UIR and UUO.

Figure S7. Gucy1α1 co-labels Pdgfrβ-, Vim- and αSma-positive fractions of kidney fibroblasts in the medullary areas under the baseline conditions and over the course of UIR and UUO.

Figure S8. Gucy1α1 expression levels increase in the prolonged UIR model of female murine CKD.

Figure S9. Myocardial infarction results in significant upregulation of Gucy1α1 along with historically used fibrosis markers Pdgfrβ-, Vim- and αSma.

Figure S10. Gucy1α1 exhibits significant correlation with αSma over the course of liver fibrosis resolution.

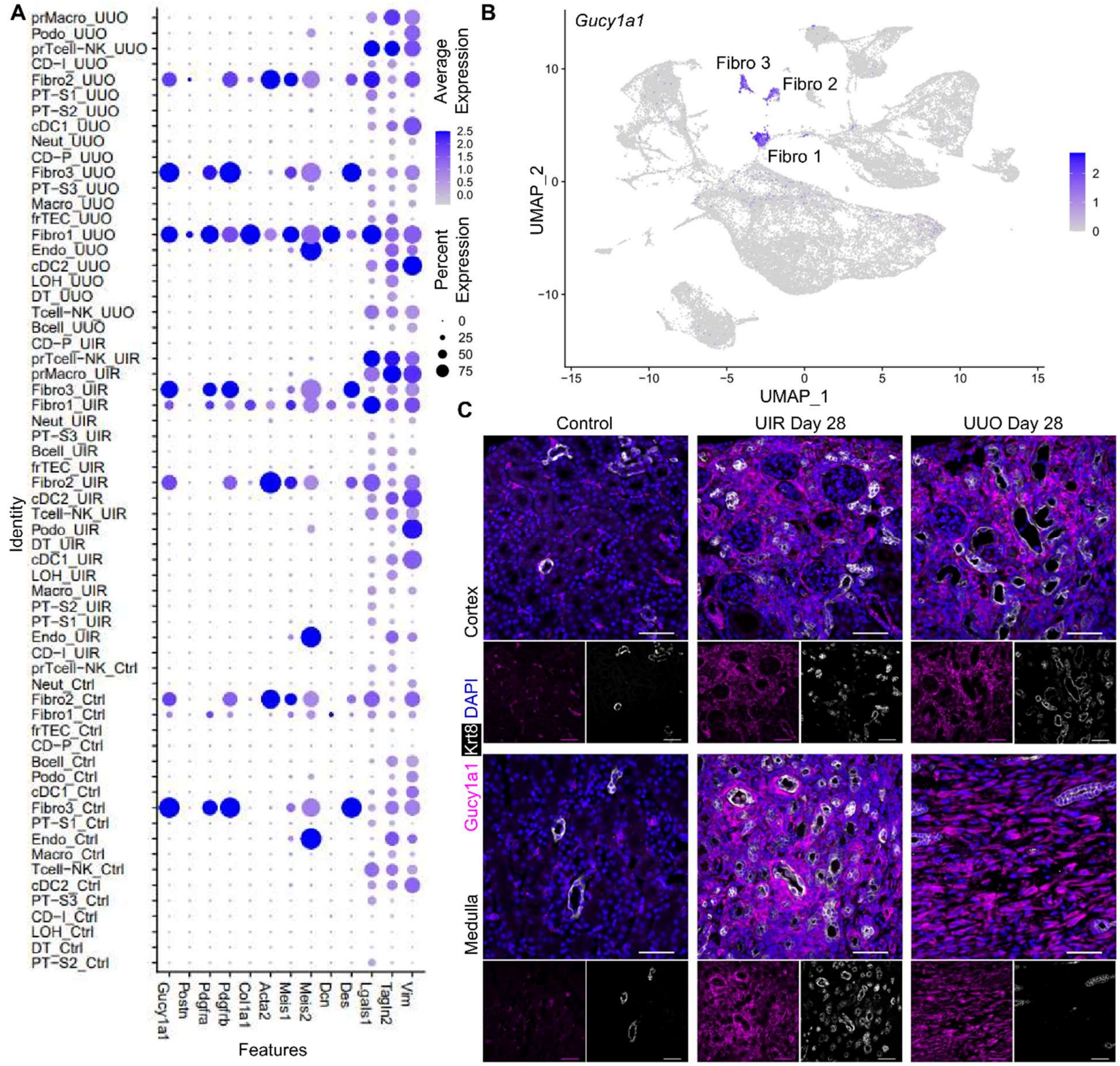


**Figure S1. *Gucy1α1* selectively marks kidney fibroblasts in two independent clinically relevant fibrosis models.** (**A**) Dot plot demonstrates *Gucy1α1* and traditionally used kidney fibroblast markers expression patterns in the control, UIR and UUO Day 28 kidneys (original dataset published in Rudman-Melnick et al., PMID: 38172172). Note that *Gucy1α1* selectively labels “Fibro 1”, “Fibro 2” and “Fibro 3” clusters of the control, UIR and UUO treated kidneys. Scale: dot size denotes percentage (0, 25, 50, 75, 100) of cells expressing the marker. Color intensity represents average gene expression values. (**B**) Feature plot demonstrating spatial patterns of *Gucy1α1* expression in the kidney cell populations. Color intensity represents average gene expression values. (**C**) IF shows Gucy1α1 expression (magenta) in the interstitial spaces of the control kidney which exhibits only mild tubular injury marker Krt8 (white) signal. Both fibrosis models elicited dramatic Gucy1α1 elevation in the interstitial intratubular cortical and medullary areas of the kidney. DAPI, blue. Original magnification, × 60, maximal intensity projection from a Z-stack, 0.28 µm/px Nyquist zoom, scale 25 µm.

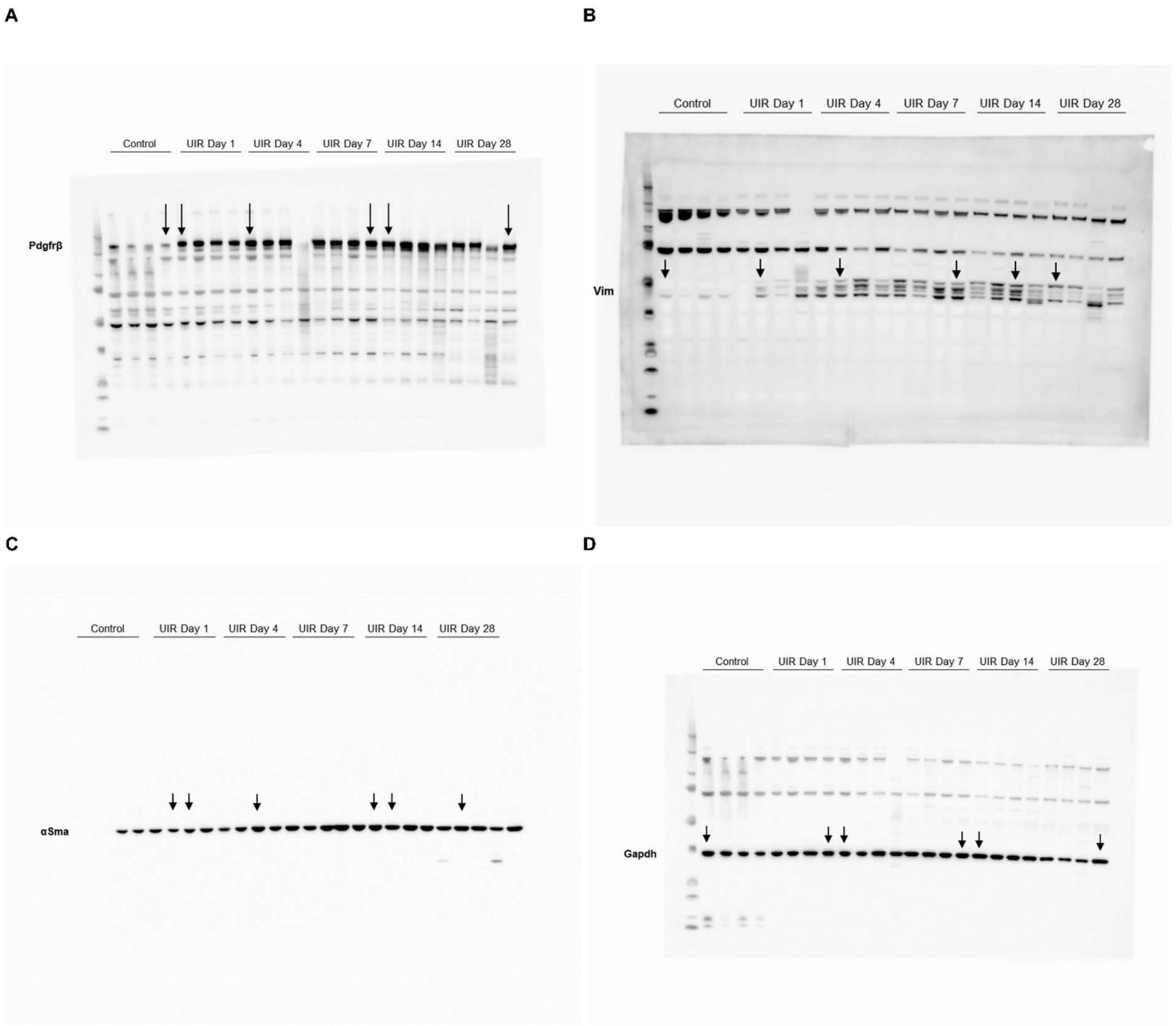


**Figure S2. UIR causes progressive elevation of fibrosis markers Vim, αSma and Pdgfrβ over the course of injury progression.** (**A-D**) UIR model of murine kidney fibrosis exhibited gradual elevation of classical stromal markers Vim (**A**), αSma (**B**) and Pdgfrβ (**C**). Gapdh (**D**) is used as loading control. Arrows are used to show representative bands demonstrated in the main Figure 1D.

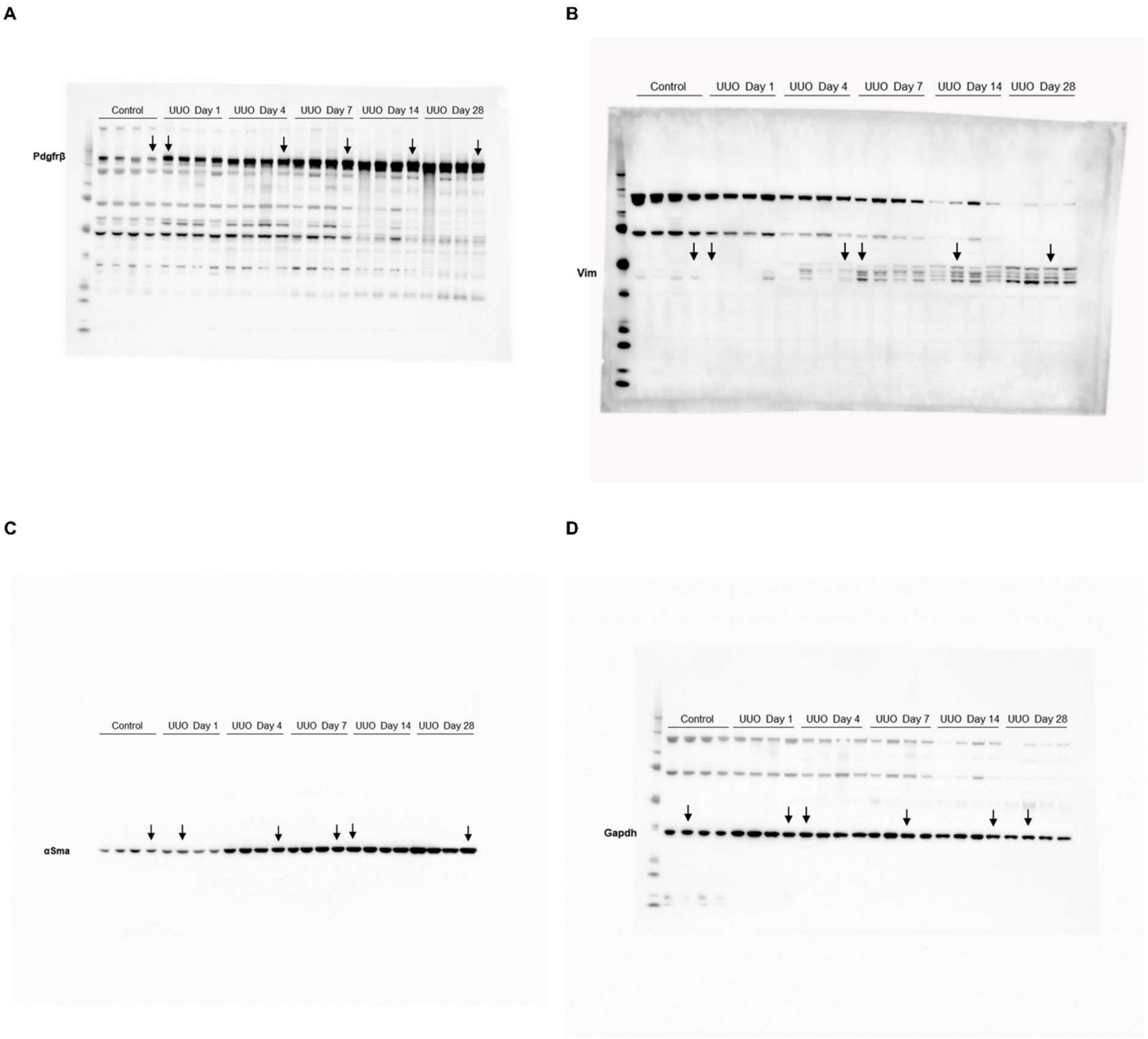


**Figure S3. UUO causes progressive elevation of fibrosis markers Vim, αSma and Pdgfrβ over the course of injury progression.** (**A-D**) UUO model of murine kidney fibrosis exhibited gradual elevation of classical stromal markers Vim (**A**), αSma (**B**) and Pdgfrβ (**C**). Gapdh (**D**) is used as loading control. Arrows are used to show representative bands demonstrated in the main Figure 1D.

A

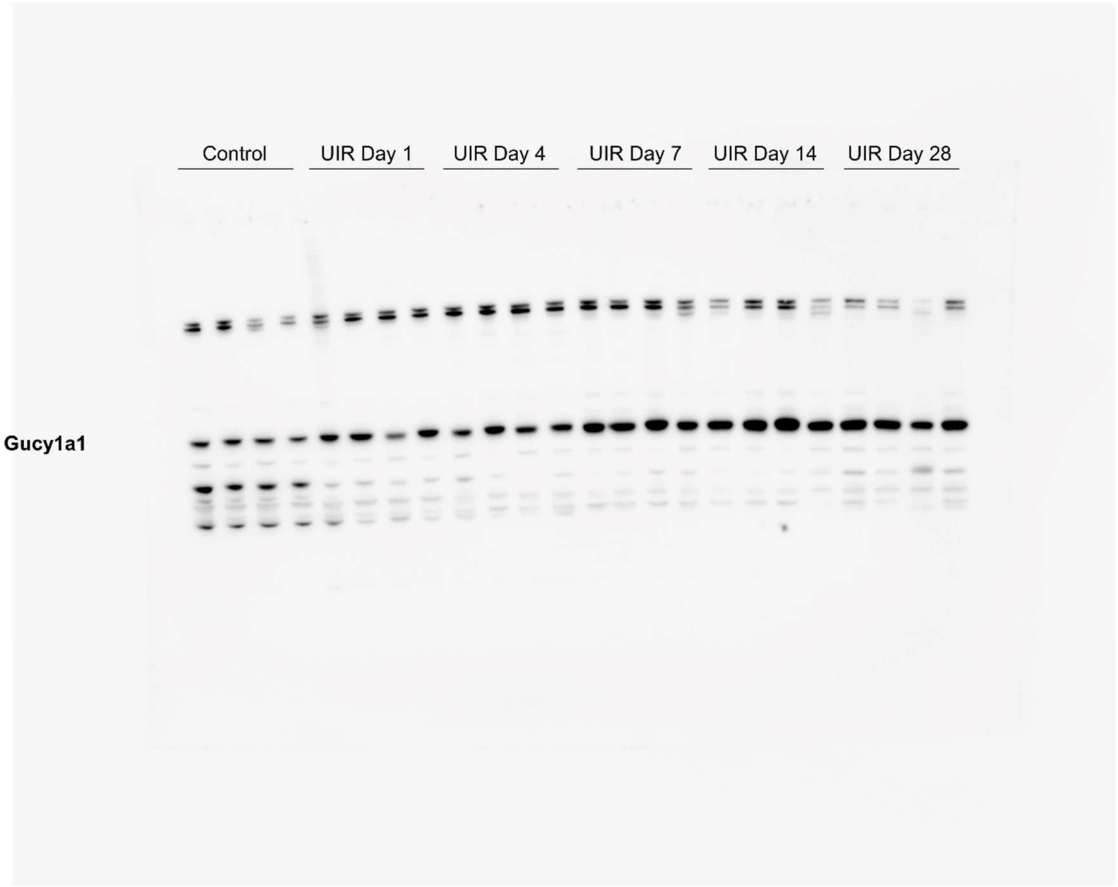


**Figure S4. UIR causes progressive elevation of Gucy1α1 over the course of injury progression.** (**A**) Western blotting demonstrates progressive changes of Gucy1α1 expression over the course of UIR induced kidney fibrosis. Mild control and early Day 1 and Day 4 expression elevates as the injury accelerates.

**A**

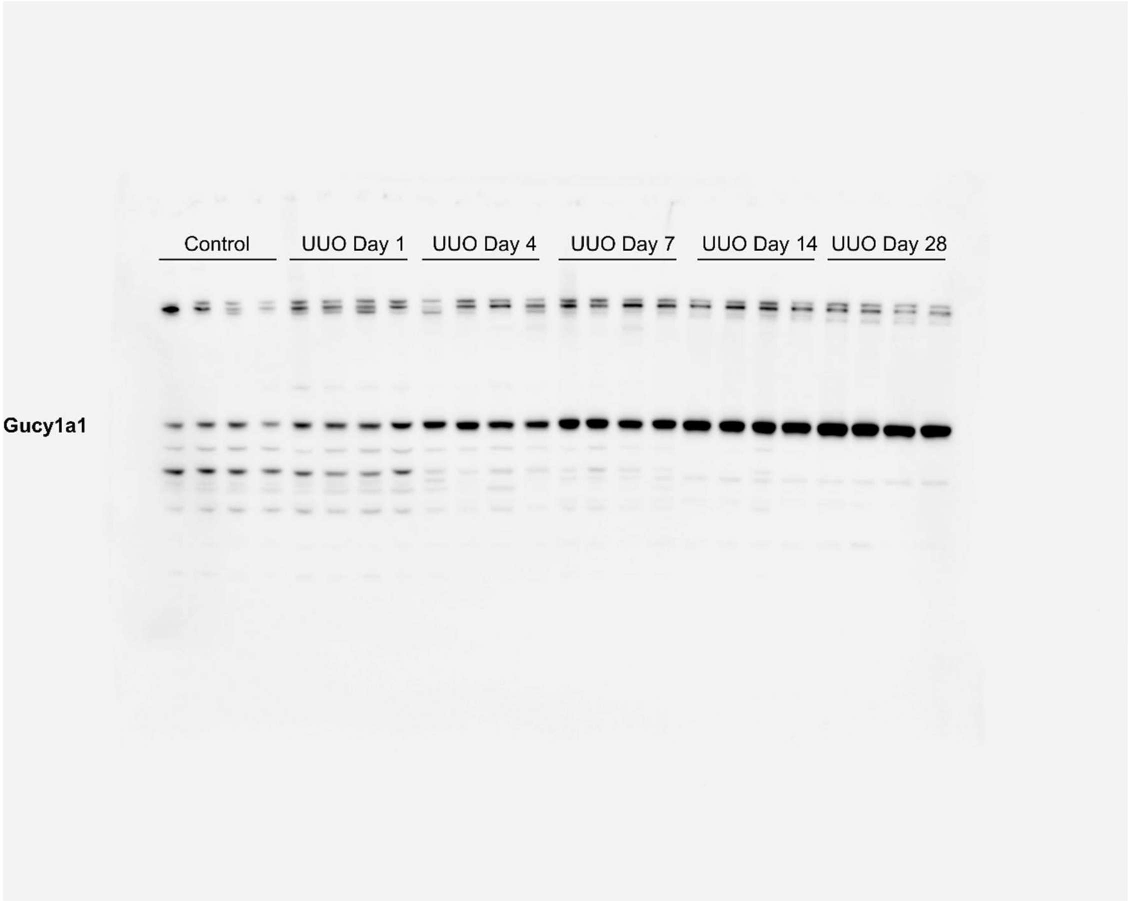


**Figure S5. UUO causes progressive elevation of Gucy1α1 over the course of injury progression.** (**A**) Western blotting demonstrates progressive changes of Gucy1α1 expression over the course of UUO induced kidney fibrosis. Note that this model causes earlier induction of fibrosis and earlier upregulation of Gucy1α1.

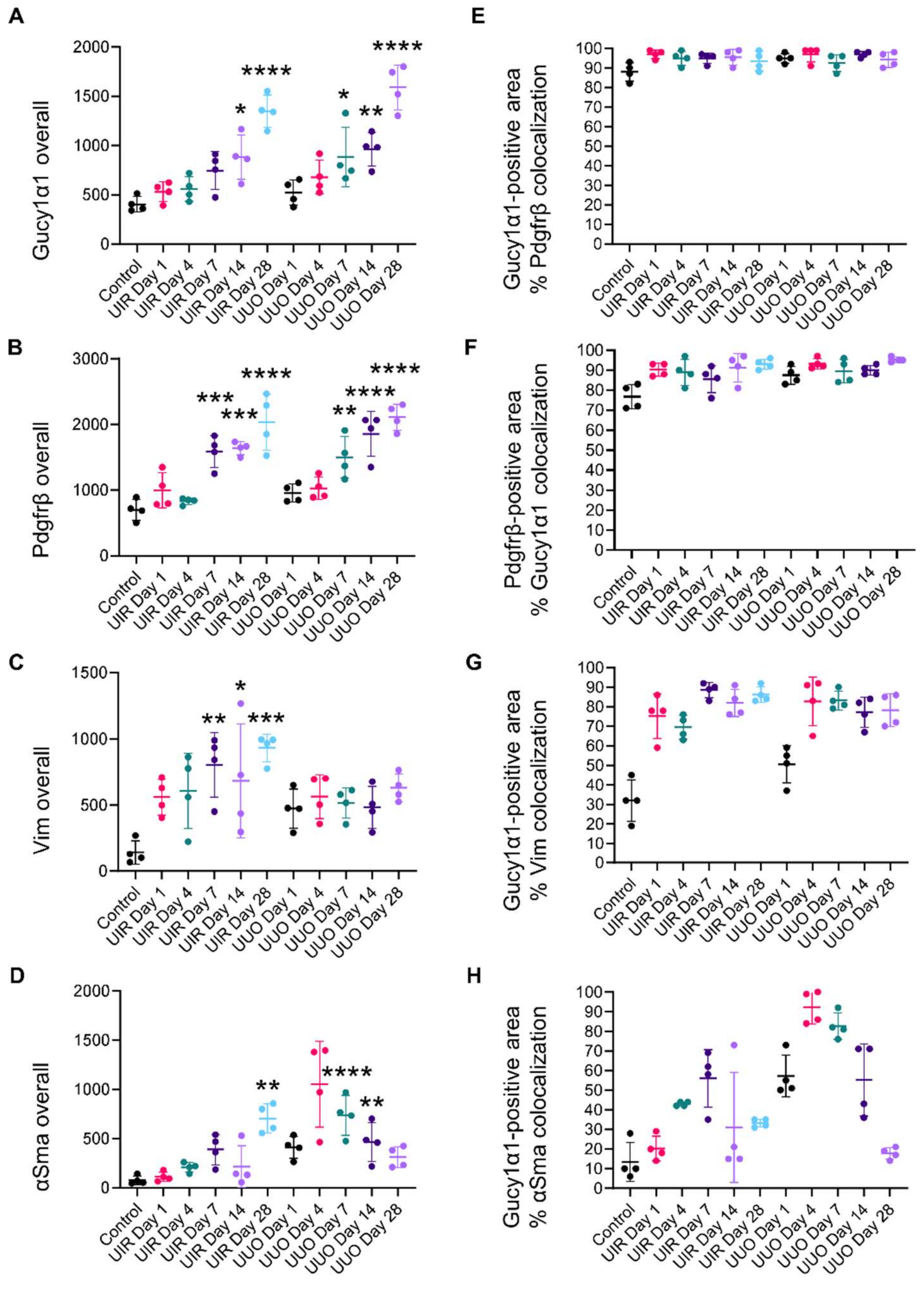


**Figure S6. Gucy1α1 co-labels Pdgfrβ-, αSma- and Vim-positive fractions of the baseline and activated kidney fibroblasts in the cortical areas.** (**A-D**) Quantitative IF analysis of overall Gucy1α1, Pdgfrβ-, αSma- and Vim levels in the control kidney and over the course of UIR/UUO progression in the cortical areas. N=4 per group, ordinary one-way ANOVA, P *≤0.05, **≤0.01, ***≤0.001, ****≤0.0001 compared to the control. (**E-H**) Scatter plots showing quantitative IF analysis of cortical patterns of kidney fibrosis markers expression. Plots show percentages of Gucy1α1-expressing cortical interstitium positivity for other fibrosis markers, including Pdgfrβ (**E**), Vim (**G**) and αSma (**H**). These graphs present data shown in Figure 3C as a violin plot. (**F**) Scatter plot showing percentages of Pdgfrβ-expressing cortical interstitium positivity for Gucy1α1. Note that not only near-total percent of Gucy1α1-expressing areas is positive for Pdgfrβ (**E**), but vice versa (percentages of Gucy1α1 co-expression among Pdgfrβ-positive cortical areas: control: 76.75%; UIR Day 1: 90.25%, Day 4: 89%, Day 7: 85.5%, Day 14: 91.25%, Day 28: 93%; UUO Day 1: 87.5%, Day 4: 93.25%, Day 7: 89.5%, Day 14: 90%, Day 28: 95.25%). N=4 animals per group. Only interstitial non-glomerular areas were included in the analysis. Data in scatter plots is shown as mean ± SD.

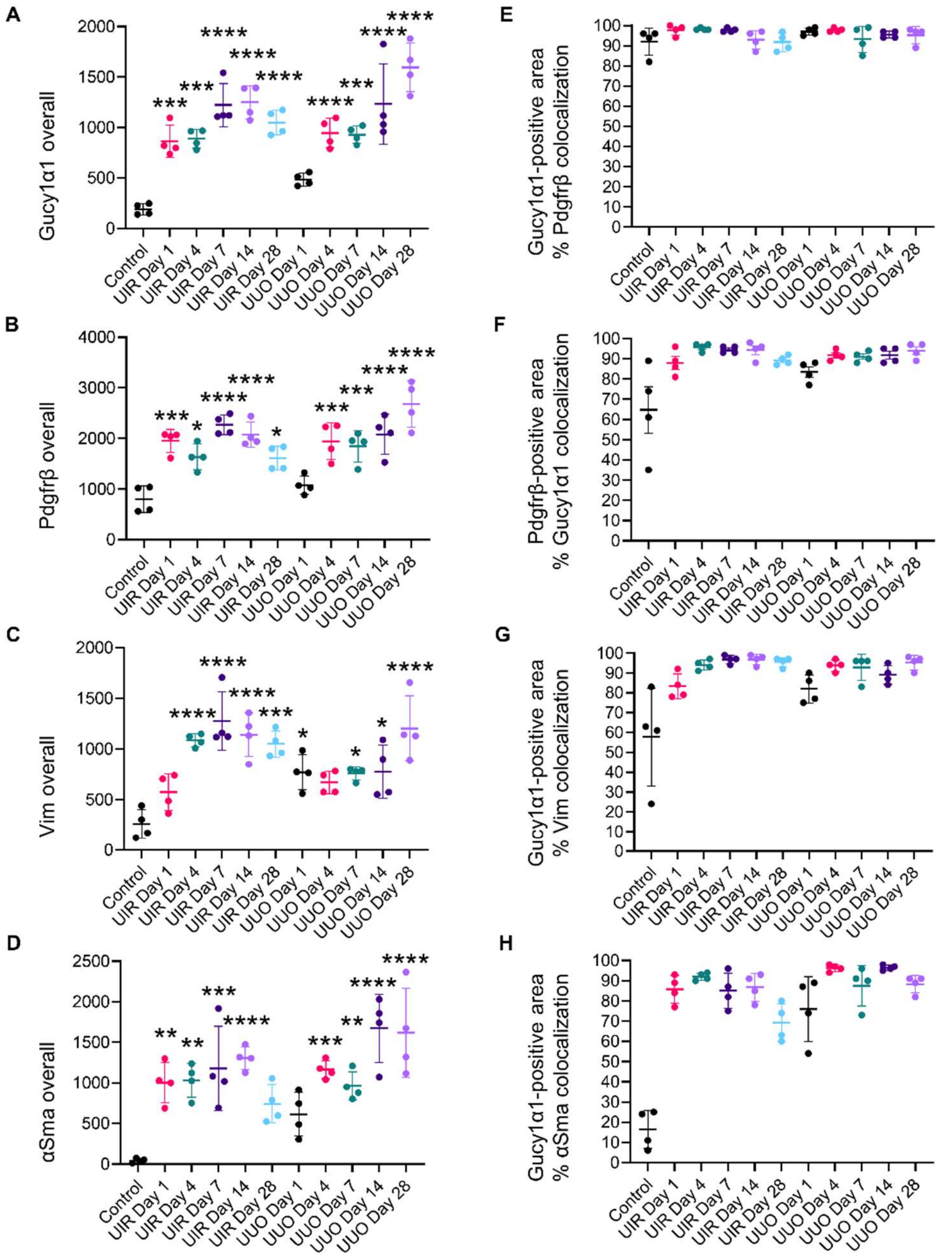


**Figure S7. Gucy1α1 co-labels Pdgfrβ-, αSma- and Vim-positive fractions of the baseline and activated kidney fibroblasts in the medullary areas.** (**A-D**) Quantitative IF analysis of overall Gucy1α1, Pdgfrβ-, αSma- and Vim levels in the control kidney and over the course of UIR/UUO progression in the medullary areas. N=4 per group, ordinary one-way ANOVA, P *≤0.05, **≤0.01, ***≤0.001, ****≤0.0001 compared to the control. (**E-H**) Scatter plots showing quantitative IF analysis of medullary patterns of kidney fibrosis markers expression. Plots show percentages of Gucy1α1-expressing medullary interstitium positivity for other fibrosis markers, including Pdgfrβ (**E**), Vim (**G**) and αSma (**H**). These graphs present data shown in Figure 4C as a violin plot. (**F**) Scatter plot showing percentages of Pdgfrβ-expressing medullary interstitium positivity for Gucy1α1. Note that not only near-total percent of Gucy1α1-expressing areas is positive for Pdgfrβ (**E**), but vice versa (percentages of Gucy1α1 co-expression among Pdgfrβ-positive medullary areas: control: 64.75%; UIR Day 1: 88%, Day 4: 97.75%, Day 7: 94.5%, Day 14: 94.25%, Day 28: 89.25%; UUO Day 1: 83.5%, Day 4: 91.75%, Day 7: 91%, Day 14: 91.75%, Day 28: 94%). N=4 animals per group. Data in scatter plots is shown as mean ± SD.

A

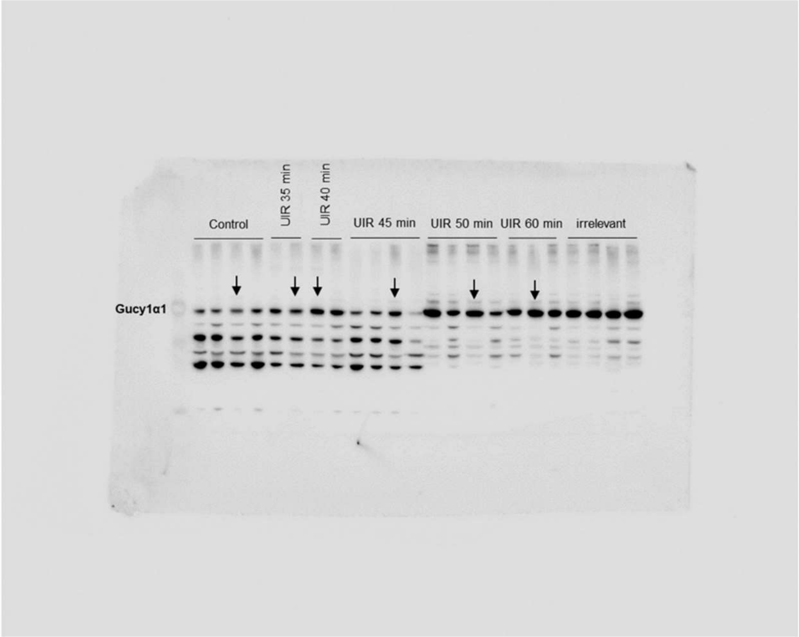


B

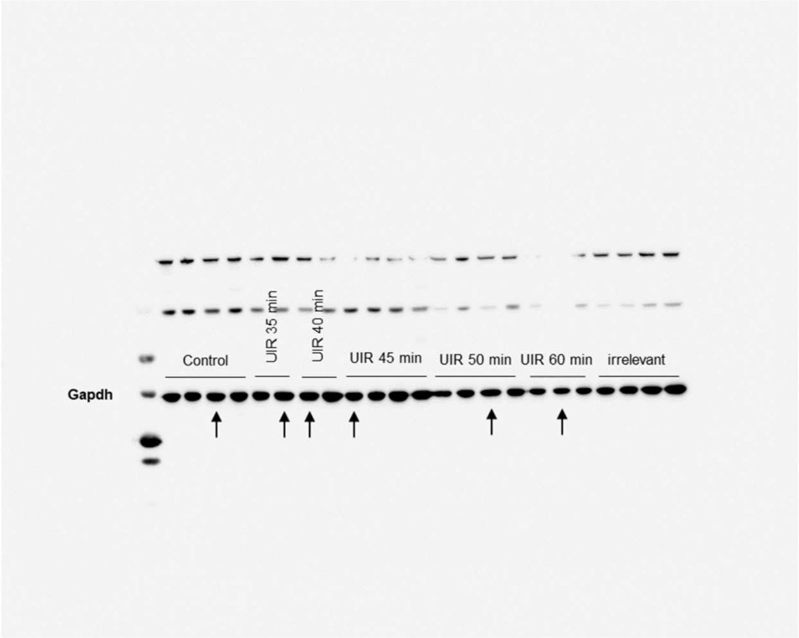


**Figure S8. UIR causes Gucy1α1 upregulation in the female model of murine CKD.** (**A** and **B**) Western blotting demonstrates changes of Gucy1α1 expression in the female control and UIR treated kidneys. Gapdh is used as a loading control. Note that prolonged UIR 50- and 60-min caused pronounced Gucy1α1 upregulation. Arrows are used to show representative bands demonstrated in the main Figure 6D.

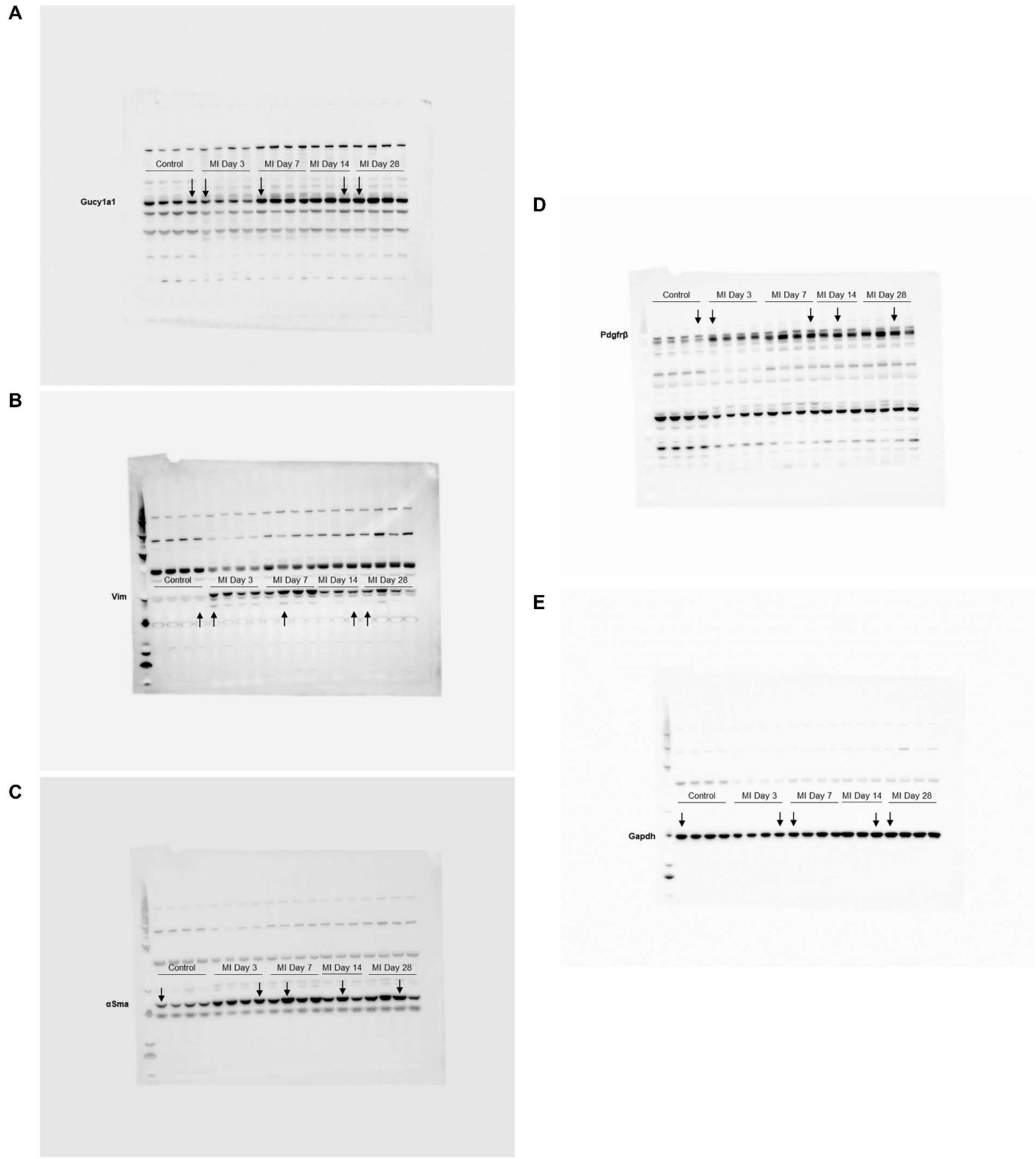


**Figure S9. Gucy1α1 labels cardiac fibroblasts in the MI model of heart fibrosis.** (**A-E**) Western blotting showing Gucy1α1, Pdgfrβ, αSma and Vim expression changes over the course of MI. Gapdh is used as a loading control. Arrows are used to show representative bands demonstrated in the main Figure 9B.

**A**

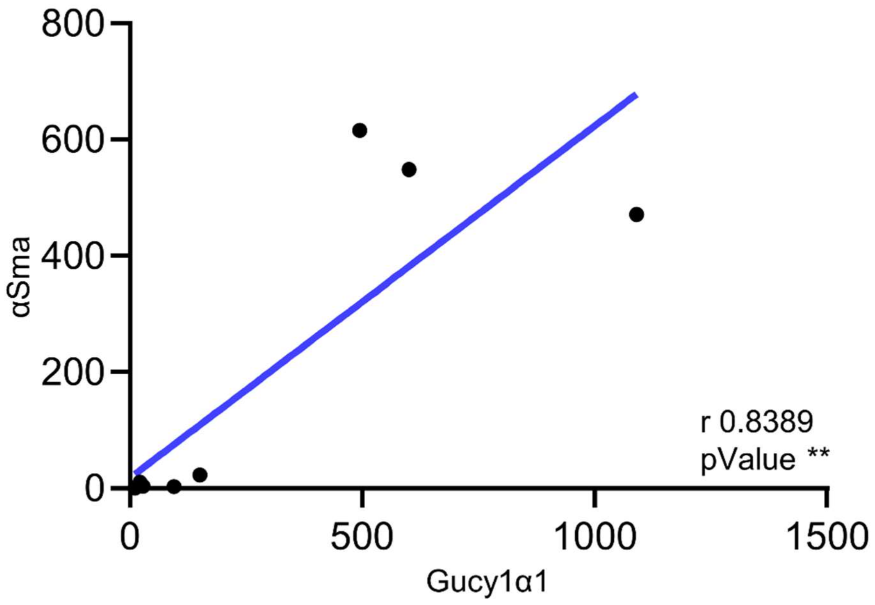


**Figure S10. Gucy1α1 directly correlates with αSma over the course of DDC induced biliary fibrosis progression.** (**A**) Correlation analysis between αSma levels and Gucy1α1 at all the timepoints detected by IF. Pearson r correlation analysis, n=9 per marker (Control, Day 14, 28, n=3 per group), r values and P as **≤0.01 for each pair are shown on the graphs presenting simple linear regression of correlation between αSma and Gucy1α1.

**Supplemental methods**

Animal procedures and harvesting. Between the experimental procedures, mice were housed with mice of the same sex at a maximum of 4 mice per cage in a specific-pathogen-free, temperature-controlled vivarium under a 12-h light–dark cycle with ad libitum access to food and water. For the surgical procedures, mice were anesthetized by 3% Isoflurane anesthesia gas before the procedures and received 1.5% Isoflurane anesthesia gas during operating. Extended release (XR) buprenorphine RODENT was administered after operating at 3.25 mg/kg body weight subcutaneously. For the organ collection, mice were anaesthetized by isoflurane inhalation and killed by exsanguination. Mice were perfused with 10-15ml of ice-cold PBS (ThermoFisher Scientific, 14190-250) via the aorta until no deep red color is observed in the kidney, then kidneys were excised, decapsulated and split in halves for snap freezing in liquid nitrogen or fixation in 4% PFA (Electron Microscopy Sciences, 15710-S) overnight (ON) at 4 °C with gentle agitation. Whole hearts were collected for fixation and histological analysis. For the molecular analysis, left ventricles were excised and snap frozen in liquid nitrogen.

Real-time quantitative PCR (RT-qPCR). Total RNA was isolated from homogenized kidney biopsies using Ambion PARIS kit (AM1921) and purified using the GeneJET RNA purification kit (ThermoFisher Scientific, KO732). cDNA was synthesized with the iScript Reverse Transcription Supermix (Bio-Rad, 1708841). qPCR was performed with TaqMan universal PCR master mix (Thermo Fisher Scientific, 4304437) on the Applied Biosystems Quant Studio 3 system with Mm012220285 *Gucy1α1* primer. The reported Ct values are the means of two cDNA sample replicates. The target gene Ct values were normalized to the eukaryotic 18S rRNA endogenous control (ThermoFisher Scientific, 4333760T) and presented as the fold change.

Western blotting. Total proteins were isolated from homogenized kidney and left ventricle biopsies using Ambion PARIS kit (AM1921). Extraction buffers were supplemented with protease (ThermoFisher Scientific, 78430) and phosphatase (SigmaAldrich, P5726, P0044) inhibitors. 10-15 µg of protein was separated via PAGE, transferred to PVDF membrane (Immobilon-FL, 0.45 µm pore size, IPFL00010), blocked with 5% non-fat milk and incubated with the target recognizing primary antibodies O/N at 4°C. Then, the membranes were washed with Tris buffered saline 0.02% Tween (TBST) and incubated with the secondary HRP-conjugated antibodies for 1 hour at RT. The target protein levels were normalized to the endogenous control detected with mouse anti-Gapdh (EMD Millipore, MAB374, 1:5000) antibody. Precision Plus protein WesternC standards (Bio-Rad, 161-0376) and Precision Protein StrepTactin-HRP conjugates (Bio-Rad, 161-0380) were used as ladder and blot detection controls. The following primary antibodies were used: rabbit anti-Gucy1α1 (Proteintech, 12605-1-AP, 1:600), rabbit anti-Pdgfrβ (Proteintech, 13449-1-AP, 1:1000), rabbit anti-Vim (Proteintech, 10366-1-AP, 1:2000), mouse anti-αSma (Sigma-Aldrich, A5228, 1:3000). The signal was visualized using SignalFire ECL reagent (Cell Signaling, #6883) using the ChemiDoc imaging system and Bio-Rad's Image Lab Touch Software. The Western blot images were analyzed using ImageJ software; the target protein signal intensity was normalized to the endogenous control protein (Gapdh) signal intensity.

Immunofluorescent (IF) staining. Fresh PFPE 6 µm kidney sections were subjected to deparaffinization, heat induced citrate epitope retrieval, permeabilization in 0.06% Triton-X-100, blocking with 4% donkey serum (Jackson Immuno-Research, 017-000-121) for

1.5 hours and O/N incubation with primary antibodies at 4°C. Primary antibodies are spun in microcentrifuge at 4°C for 10min at 14000 rpm. Following primary antibodies were used for IF: rabbit anti-Gucy1α1 (Proteintech, 12605-1-AP, 1:75), goat anti-Pdgfrβ (R&D Systems, AF1042, 1:40), mouse anti-αSma (Sigma-Aldrich, C6198, 1:400), guinea pig anti-Nephrin (Progen, GP-N2, 1:50), rat anti-Krt8 (EMD Millipore, MABT329, 1:100), chicken anti-Vim (EMD Millipore, AB5733, 1:150). Then sections were incubated with secondary fluorescent antibodies, stained with DAPI (ThermoFisher Scientific, 62247) and mounted with Vectashield antifade mounting medium (Vector Laboratories, H-1000-10). The following secondary antibodies from Jackson Immuno-Research were used: Alexa Fluor 647-conjugated AffiniPure donkey anti-rabbit, 711-605-152, 1:500; Alexa Fluor 488-conjugated AffiniPure donkey anti-chicken, 703-545-155, 1:500; Alexa Fluor 594-conjugated AffiniPure donkey anti-guinea pig, 706-585-148, 1:100; Alexa Fluor 594-conjugated AffiniPure donkey anti-rat, 712-585-153, 1:100; Alexa Fluor 594-conjugated AffiniPure donkey anti-goat, ab175745, 1:250. All secondary antibodies were spun before and after the dilution in 1% donkey serum in microcentrifuge at 4°C for 15min at 14000 rpm.

<u>Picrosirius Red staining</u>. PFPE 6 μm kidney sections were rehydrated in a series of EtOH dilutions, incubated with Picrosirius Red solution (Abcam, ab246832) for 1 hour, rinsed with glacial acetic acid aqueous solution and mounted with Permount medium (ThermoFisher Scientific, SP15-100).

<u>Microscopy and image analysis</u>. For quantitative IF analysis, ×60 0.28 μm/px resolution Z-stack images were obtained. For overall signal analysis, Imaris algorithm "Spots" was used to detect the target protein expression, and the total spot number detected in the

visual field was reported. For the analysis of intraglomerular Gucy1α1, Pdgfrβ and Vim expression, Imaris algorithm “Surfaces” was used to indicate Nphs1-expressing podocytes. Then, “Spots” were used to detect Gucy1α1, Pdgfrβ and Vim signal inside Nphs1-positive surfaces. The number of detected spots was normalized to the surface volume and averaged among all glomeruli captured per visual field. For the analysis of Gucy1α1-expressing area colocalization with other fibrosis markers, Imaris algorithm “Surfaces” and spatial colocalization were used. Channels showing Gucy1α1, Pdgfrβ, αSma and Vim signals were labeled with corresponding “Spots”.

To determine the percentage of Gucy1α1-expressing areas colocalizing with other markers, those spots located within 10μm from Gucy1α1 spots were counted in as a colocalized fraction. Vice versa, to determine the percentage of Pdgfrβ-positive areal also positive for Gucy1α1, those Gucy1α1 spots located within 10μm from Pdgfrβ spots were counted in. Identical “Spot” and “Surface” settings were used for all the conditions across experimental groups. For the cortical areas analysis, all Gucy1α1, Pdgfrβ, αSma and Vim spots located inside Nphs1-positive areas were excluded from analysis.

<u>Schematic.</u> Heart and kidney schematic images from Figure 1A, 3A, 4A and 9A were created in Illustrator. Liver schematic images from Figure 10A were created in BioRender.

<u>Data availability</u>. scRNA-seq data referred in the manuscript is deposited at the Gene Expression Omnibus under accession number GSE198621.
